# Missense mutation knowledge can decrease prediction inaccuracies on protein secondary structure

**DOI:** 10.1101/2025.03.04.641402

**Authors:** Ivan Perez, Ulrike Stege, Hosna Jabbari

## Abstract

Protein tertiary structure prediction models like AlphaFold2 have revolutionized the field with unprecedented accuracy. Yet predicting structural changes arising from single amino acid mutations remains a challenge. The complexity introduced by these mutations calls for models that can incorporate mutational information into their predictions. We propose a novel refinement strategy for protein secondary structure prediction that leverages missense mutational data. As part of this strategy, we introduce *Mut2Dens*, a model that not only yields improved consistency of predictions for mutational data, but also maintains robust predictive performance on non-mutational datasets. Mut2Dens takes multiple predicted secondary structures and generates a mutation-aware secondary structure. This awareness comes from our mutational dataset, learning to avoid common mistakes in prediction methods after a missense mutation occurs. In particular, Mut2Dens employs the extremely randomized trees (ExtraTree) algorithm to avoid overfitting and makes effective use of the limited mutational data available from experimentally determined three-dimensional structures. By combining predictions from highly accurate structure prediction models, we create an ensemble that integrates their strengths while enhancing mutational capabilities. This refinement strategy also improves the non-mutational performance of state-of-the-art methods by addressing their most inaccurate and least confident predictions. Moreover, it reduces improbable outcomes in mutated protein structures—such as transforming *π*-helices into *β*-sheets—that can still occur in current prediction models. Finally, by using interpretable machine learning algorithms (e.g., ExtraTree), we can reveal the underlying biological knowledge from the refinement model; the insights gained from Mut2Dens can be corroborated with known mutational outcomes, helping users pinpoint discrepancies across structure prediction models and make more informed decisions regarding the predicted structures. The data utilized here is available at https://github.com/ivanpmartell/sam-models.

## Introduction

The three-dimensional structure of a protein is directly correlated with its function and is partly defined by its amino acid sequence [1]. Recently, protein tertiary structure prediction models have become highly accurate [2] for proteins with known homology. Despite these achievements, it is an open question on whether these models are able to correctly predict structural outcomes caused by single amino acid mutations [3, 4].

Protein three-dimensional structure is inherently noisy due to the dynamic nature of atomic positions. This includes atomic vibrations, environmental conditions during experimentation, and the inherent flexibility of protein structures; therefore, making it challenging to obtain a precise and stable representation of the protein’s architecture. Protein secondary structure helps mitigate these inconsistencies and variations introduced by noisy atomic coordinates and atomic-level fluctuations by discretizing the atomic-level details, thereby elucidating mutational effects in the protein’s architecture or backbone structure. Specifically, the distances between backbone atoms are used to infer backbone bonds, which can then be classified into secondary structure elements (SSEs) by secondary structure assignment algorithms such as DSSP [5]. This removal of confounding atomic-level variations introduced by experimental and physical factors can increase the reliability of observed structural changes.

Recent studies [3, 4] evaluated highly performing structure prediction models on missense mutations. These evaluations, however, only focused on backbone information that utilize *three-class secondary structure* (Q3). As the name implies, Q3 classifies all SSE into three classes characterized by their secondary structure—*α*-helix, *β*-sheet, and coil. However, as documented in the literature [5, 6], Q3 is insufficient to account for the complete structural information of the protein backbone. Moreover, as existing methods for predicting secondary structures near the theoretical accuracy limit of nearly 90% for three-state predictions [7], attention has turned to the more complex challenge of predicting *eight-class secondary structure* (Q8). While the theoretical limit of Q8 accuracy is not well established, current template-less methods achieve an accuracy of about 75% [8].

Evaluating secondary structure prediction requires a consistent secondary structure assignment to serve as a *gold standard*. However, most secondary structure assignment algorithms produce varying results [9], especially in proteins with irregular conformations [10]. We selected DSSP [5], a pioneering algorithm for secondary structure assignment, as the ground truth for machine learning (ML) purposes because it has been extensively tested and widely used.

A previous investigation into the secondary structure of predicted proteins, intended to gauge how well state-of-the-art models account for structural effects from mutations, found inconsistent predictions for altered secondary structure elements [11]. Although the overall predicted secondary structure remains accurate, these models often locate mutational changes incorrectly, rendering them unsuitable for mutational prediction. Therefore, we propose a refinement strategy for protein secondary structure prediction that enhances already accurate models for mutational data and further benefits their non-mutational predictions. This approach utilizes an ensemble model, Mut2Dens, which is trained on missense mutation data, and increases the prediction scores of secondary structures by at least 20% in low-scoring proteins. Moreover, Mut2Dens demonstrates substantial improvements in maintaining mutational consistency for altered secondary structure elements.

## Background

Protein structure is described through multiple differing levels of complexity. *Primary structure* refers to the linear sequence of amino acids, linked by peptide bonds, which make up the protein chain and dictates all subsequent levels of its structure. *Secondary structure* refers to the conformation of the protein backbone excluding the side chains. Regularly occurring secondary structures in proteins are called elements, e.g. *α*-helices and *β*-sheets. These secondary structure elements can vary widely in length, from as few as three to five residues, to over fifty residues. The connectivity between such secondary structure elements is often referred to as the protein topology. *Tertiary structure* is the overall three-dimensional shape or fold of a protein chain, stabilized by physico-chemical interactions to its surrounding environment, and thus heavily influenced by it. Lastly, the *quaternary structure* is the arrangement of multiple folded protein molecules in a multi-subunit complex.

Protein tertiary structure dictates the protein’s overall shape with the three-dimensional arrangement of all its atoms relating to the precise spatial coordination of secondary structure elements. Since tertiary structure highly depends on a dynamic environment, its shape is constantly fluctuating, albeit with some stability from inter-atomic bonds. This is corroborated by the hierarchic model of protein folding, which dictates that pre-organized elements of local secondary structure fold successively into ever-larger superstructures with native-like topology. Therefore, protein backbone and its secondary structure elements and overall topology can be utilized to detect folding transitions and changes to the protein shape without the need for atomic level coordinates that can become chaotic from subtle changes, e.g. Brownian motion and differing experimental conditions.

Since the inception of secondary-structure prediction models, evaluations have relied on accuracy and segment-overlap metrics [12, 13], which have been continuously refined over the years [14]. Although these metrics are valuable for assessing secondary-structure predictions, further work is needed to adapt them for evaluating mutational structural changes. To measure predictions from mutational results, we utilize previously devised mutational metrics used while benchmarking structure prediction models on single amino acid mutations [11]. More specifically, we focus on mutational consistency that can distinguish the performance between disruptive and stable structural changes that occur due to a mutation. When a mutation induces a structural change, we refer to it as *disruptive*. Therefore, an SSE has changed due to an amino acid mutation. In contrast, if no SSE change occurs, the mutation is considered *stable*.

There are multiple types of models that can produce accurate protein structure predictions. The models differ by their resulting level of structure complexity: secondary and tertiary structure prediction models. We explore both types, as both can produce an output that culminates in a secondary structure prediction.

As the review by Ho et al. [15] points out, while the predictive performances of recent secondary structure prediction models are similar to each other, differences arise when predicting secondary structures based on Q8 assignment. Therefore, we carefully selected publicly available state-of-the-art protein structure prediction models for consideration into our ensemble. These considered models include Raptor-X Predict Property tool [16], SSPro8 [6], SPOT-1D [17], SPOT-1D-Single [18], and SPOT-1D-LM [19]. We excluded models that are currently unavailable or required extensive training, in the case of deep learning models without publicly available pre-trained models. Furthermore, we solely procure protein structure prediction software that can be installed on-premise. This decision is based on our observation that cloud-based software are often no longer supported after a few years. PsiPred, although one of the pioneering secondary structure prediction models, was excluded as it is solely a Q3 predictor. Similar to PsiPred, other Q3 prediction software that were not included in this work consist of PHD [20], SOPMA [21], SPINE-X [22], SPARROW [23], and JPRED [24]. Other Q8 prediction software that were disregarded include SPIDER3 [25] and SPIDER3-Single [26], as SPOT-1D and SPOT-1D-Single have superseded these tools.

Tertiary structure prediction methods were analyzed and selected for their exceptional results in predicting tertiary structure, as well as their contrasting prediction procedures albeit their consistent use of deep learning algorithms. Deep learning algorithms pioneered by AlphaFold2 [2] have enabled unprecedented progress by extrapolating the three-dimensional structure from a protein’s homologs through its shared ancestry. For AlphaFold2 and its derivatives [27], e.g. ColabFold [28], homology is obtained using sequence similarity — performed through multiple sequence alignment (MSA) of the target protein to other known protein sequences. For language models such as ESMFold [29, 30] and RGN2 [31], homology is obtained while training the protein language model and is included in the trained parameters of the model. The previously mentioned list of tertiary structure prediction methods — Alphafold2, Colabfold, ESMFold, and RGN2 — along the considered secondary structure prediction methods listed above make up the complete list of considered predictors for the ensemble.

An ensemble of these protein structure prediction models requires computational methods capable of processing the information of each predictor and refine them into an consolidated result. A naive method would include a majority algorithm that selects the most commonly predicted Q8 element for each amino acid in the sequence. Machine learning (ML) methods expand this idea by producing a weighted selection depending on each predictor’s outcome. Recently, deep learning has become the leading machine learning methodology for prediction performance, yielding an increase by its use in many fields of research [15, 32, 33]. Due to challenges in understanding how deep learning models generate their predictions, additional investigation into the capabilities and limitations of these models is necessary [34, 35]. Therefore, interpretable machine learning methods are favored when their performance is comparable to deep learning models, as it is easier to elucidate biological meaning from the latter.

With the advances of highly accurate predictive models, machine learning models need to be understood and trusted to perform their assigned tasks [36]. In the natural sciences, interpretable models can give researchers the knowledge required to make informed decisions about the benefits and consequences arising from the predicted outcome. For this reason, we opted to investigate both deep learning models and highly interpretable ML models, e.g. decision trees.

Previous studies [37, 11] have shown that current protein structure prediction models have not effectively incorporated mutational outcomes into their predictions. In the coming sections, we introduce our novel refinement strategy, leveraging mutational data to enhance the structure predictions that occur from single amino acid mutations. Our strategy is materialized in Mut2Dens, an interpretable ensemble model specifically designed to integrate missense mutation information through our refinement strategy. Mut2Dens improves predictions for mutational data while maintaining robust performance on non-mutational datasets. By combining the strengths of multiple predictors, our ensemble model significantly enhances predictive reliability, reducing inaccuracies and improbable outcomes in mutated protein structures. We evaluate Mut2Dens in mutational and non-mutational data and analyze the interpretable model to obtain biological knowledge gathered from its training.

## Machine learning algorithms

Our refinement strategy requires the creation of a machine learning model that will transform the outputs from all structure predictors into a finalized secondary structure prediction. There exists a myriad of machine learning algorithms to create such a model. We investigate both parametric and nonparametric types of supervised ML algorithms for this task. We categorize these algorithms into:

1. *Tree*-type ML algorithms, which create a tree-like structure of optimal decision splits to determine an outcome.
2. *Neural*-type ML algorithms, which create an interconnected network-like structure of linear and non-linear functions.

For our investigation into parametric approaches, we selected neural-type algorithms for their advances and continuous improvement in performance across various fields of study. As for the nonparametric methods, tree-type algorithms were selected for their high interpretability and their ability to capture non-linear relationships without the need of feature scaling. Despite the availability of numerous other machine learning algorithms [38], we confine our analysis to the algorithms previously described to corroborate our refinement strategy for missense mutational data. Our refinement strategy aims to improve current prediction models that typically exclude or restrict protein sequences with high similarity or sequence identity during their training procedure.

The investigated tree-type algorithms include decision trees [39], random forests [40], and ExtraTree [41]. A decision tree works by splitting the data into subsets based on the value of input features. This process is repeated recursively, creating a tree-like structure where each internal node represents a decision based on a feature, each branch represents the outcome of the decision, and each leaf node represents a possible outcome. Decision trees are easy to interpret and visualize but can be prone to overfitting [42].

A random forest model consists of multiple decision trees during training and outputs a majority prediction from all trees. It introduces randomness by selecting a random subset of features for each tree and using bootstrap samples of the data. ExtraTree models are similar to random forests but introduce more randomness to the decision splits. At each split, the decision threshold is drawn at random for each candidate feature and the best of these randomly-generated thresholds is selected. This results in a more diverse set of trees and can lead to a reduction in variance [41]. ExtraTree models are computationally efficient and can handle large datasets effectively. The randomization also reduces overfitting and improves generalization. A depiction of the tree-type algorithms can be seen in Fig. 1.

**Figure 1.**
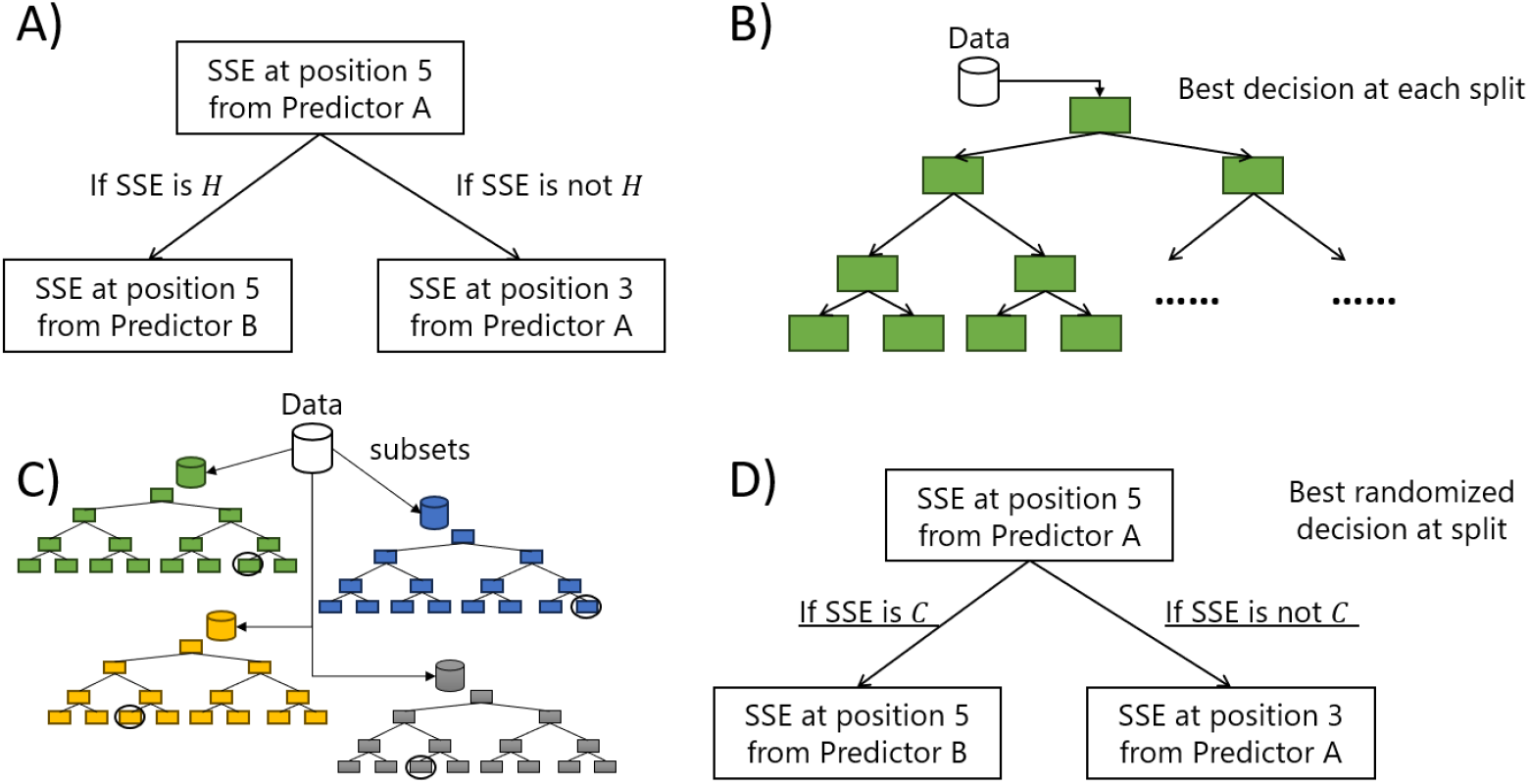
Tree algorithms. Summary of tree-type machine learning algorithms. A) Depiction of a decision threshold for tree-type algorithms. Squares contain the threshold while the arrows show possible decisions depending on the input. A)Depiction of a Decision tree model, which has a tree-like (directed acyclic) graph. The best decision thresholds are created according to a quality criteria (Gini impurity). All data is utilized to create the tree. C) Random forest model depiction showing the data being split and used to create multiple decision trees. A majority vote will become the final decision of the trees. D) ExtraTree model depiction, which functions similarly to a Random Forest but where decision thresholds are randomly selected and the best random threshold is utilized.

Neural-type algorithms create models that consist of layers of interconnected nodes, or neurons, which process its input data and passes that processed input to other neurons depending on activation functions in the network. Each neuron processes its input by utilizing learnable parameters in linear functions that get optimized to match the desired output through gradient descent. The activation functions introduce non-linearity, which allows the modeling of complex patterns of data. Architectures of the neural-type algorithms investigated include fully-connected, convolutional, recurrent, and transformer architectures [43].

Fully connected neural networks [44] consist of neuron layers where each neuron is connected to every neuron in the previous and next layers. Convolutional neural networks (CNN) [45] are designed to process grid-like data, such as images. They consist of convolutional layers of neurons where learnable parameters are reused to capture local patterns in the data. Recurrent neural networks (RNN) [46] have neurons that form directed cycles which reuse learnable parameters to maintain a “memory” of previous inputs. This makes RNNs suitable for tasks like language modeling and sequence prediction. Transformer neural networks are also designed to handle sequential data but differ from RNNs by using self-attention mechanisms [47] to process all elements of the sequence simultaneously.

### Feature selection methods

The prediction of protein secondary structure can take a considerable amount of time depending on the predictor. As our refinement model transform the outputs from selected structure predictors into a finalized secondary structure prediction, each predictor’s outcome must be computed before the refinement process can commence. From an environmentally-conscious computational performance perspective, we narrow the list of considered structure predictors through feature selection procedures. This limitation ensures a refinement model that utilizes the most important structure predictors, while removing the environmental footprint from the least important structure predictors. The computational performance, and thus footprint of each predictor is shown in Section 4 of the Supplementary materials. We utilized the following three different statistical methods to confirm the importance of each predictor.

- *χ*^2^ [48]: Compares the observed frequencies in each category to the frequencies expected. A greater difference between the observed and expected frequencies suggest a potential association or significance from the input to the predicted outcome.
- Mutual Information [49]: Quantifies the amount of dependence between two variables. Greater values indicate higher certainty of the predicted outcome by the input.
- Analysis of Variance (ANOVA) [48]: Compares the mean and variance between and within groups to determine the statistical significance between the groups. This process identifies the amount of influence that groups have on the predicted outcome.

## Materials and methods

Our refinement strategy utilizes the predicted outcomes of already established structure prediction methods, also known as predictors. The predictors’ outputs are combined and used as input to a machine learning model that refines the predictions into a consolidated result. This model is also known as an ensemble model for its ability to combine multiple predictions and refine them for mutational purposes. In this section, we describe the data utilized for training the ensemble model. We also describe the testing datasets used to validate models from the refinement strategy, and compare them to the individual predictors. We also describe the metrics used for the validation process of secondary structures, mutational changes, and to quantify the spread of the predictions from models.

### Data acquisition and pre-processing

A mutational training dataset and two test datasets were utilized in this study. The mutational training dataset consists of clusters of mutated proteins containing experimental data from the Protein Data Bank [50] (PDB). Protein sequences (PDB_SEQRES.TXT) and their experimentally derived three-dimensional structures (PROTEIN.CIF) were obtained from the PDB as of April 2023. We excluded non-protein sequences and duplicates, retaining only proteins composed of the standard 20 amino acids. Consequently, any sequences containing ambiguous amino acids were also removed.

We clustered the protein sequences using CD-HIT [51] with a 99% sequence similarity threshold, producing groups of mutated proteins. Within each cluster, we performed multiple sequence alignments using Clustal Omega [52] to identify amino acid substitutions and discard incomplete sequences. After alignment and filtering, each cluster retained only full-length sequences of uniform length.

However, while retrieving the structural data files for these sequences, we occasionally encountered structures with missing amino acids, resulting in gaps where the atoms’ locations are inconclusive. Such gaps can obscure the true effects of missense mutations, as the three-dimensional structures may not fully represent the corresponding protein sequence associated to the structure. To ensure that all relevant atomic positions are accounted for, we excluded any protein whose structure file contained such gaps. This approach guarantees that the mutation effects that we analyze are accurately represented.

After this pre-processing step, we applied DSSP to assign secondary structures to each protein sequence using their corresponding experimental structures. As resulting protein sequences had uniform length within a cluster and differ through missense mutations, the proteins could be considered aligned. Wild-type and mutated sequences were identified through mutation extraction that is detailed in Section 1 of the Supplementary materials.

We ran each method on our dataset and normalized the results to ensure consistent Q8 predictions, as different methods may use distinct symbols for identical secondary structure classes. After completing these steps, we obtained the final preprocessed dataset used for evaluating the methods. More information on the mutational dataset and its statistics can be found in a previous study [11].

Our testing datasets include the commonly utilized CB513 dataset [53] used for evaluation of secondary structure prediction models. Unfortunately, since this dataset has been extensively used, we cannot guarantee that predictors have not been trained on this data. Additionally, we obtained data from CASP15 [54] to avoid any potential overlap with training data.

The CB513 dataset is obtained from the creators of JPred (https://www.compbio.dundee.ac.uk/jpred/downloads/513_distribute.tar.gz). Additionally, we filter this dataset by remove sequences shorter than 30 amino acids, as well as any sequences containing ambiguous amino acids. We created the CASP15 dataset by obtaining the three-dimensional experimental data from the target list of regular proteins from the 15^th^ iteration of the Critical Assessment of Structure Prediction (CASP) competition. This data is hosted in the PDB and was released after the conclusion of 15^th^ CASP competition (https://predictioncenter.org/casp15/). For the CASP15 dataset, we filtered protein sequences that were too long, greater than 1024 amino acids, to accommodate predictor’s sequence length limitations.

### Input data

We investigated multiple ways in which to represent the data used as input to our ensemble model. These input representations differ due to the requirements of the type of machine learning algorithm. For neural-type, the data was organized into a three-dimensional (*i* × *j* × *k*)-matrix containing one-hot encoded vectors. Here, *i* denotes the number of secondary structure classes (which is eight for Q8 prediction), *j* denotes the sequence length and *k* denotes the number of predictors.

Machine learning frameworks for designing tree-type algorithms, e.g. Scikit-learn [55], require each sample to be a one-dimensional vector to ensure consistency of the feature representation where each element in the vector corresponds to a single feature from the sample. Therefore, tree-type algorithms were limited to a one-dimensional vector of features which could be organized into different representations. We represented sample vectors in the following manners:

- Nominal data: (*n* ·*m*)-vector, where *n* denotes the sequence length and *m* denotes the number of predictors. See Fig. 2 C for an example of this representation with 3 predictors and a sequence length of 9.
- Windowed nominal data: Same as nominal data but the sequence is truncated to a specific window size. The full sequence is processed through a sliding window with a step size of 1, where the centre location of the window has the amino acid of interest. See Fig. 2 D for an example containing a window size of 7.

**Figure 2.**
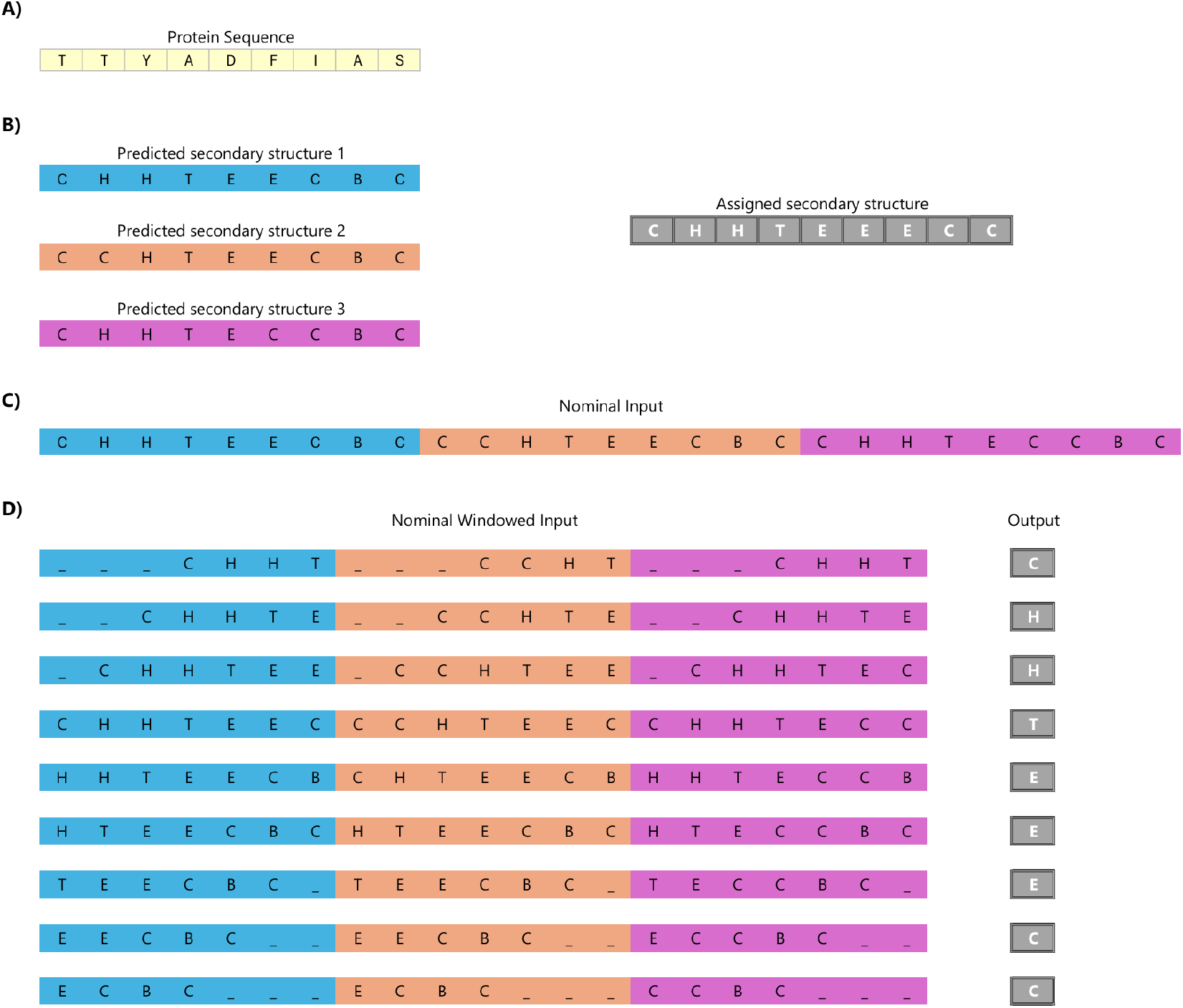
Data representation. Secondary structure representation as input features for machine learning and feature selection procedures. A) Example of a protein sequence of length 9. B) Output from three structure predictors and the assigned secondary structure to the protein sequence. C) Nominal data representation. This representation concatenates all predictions with their full length. D) Windowed nominal data representation. Window size of 7. To differentiate parts of the sequence, the data is padded with empty spaces. Each row represents an input with a position of interest. This position is located in the middle of the window, and also contains the SSE output.

Because the input sequence length can vary, we use a window-based approach that allows the user to specify the maximum length. We tested prime numbers (starting at 3) for the window lengths as the window sizes need to be odd to place the amino acid of interest at the center. Prime numbers were specifically selected to follow the distribution from the prime number theorem [56] — prime numbers become less common as they become larger. Window sizes in our data seem to affect outcomes more sensitively the shorter they were, and thus we surveyed shorter sized windows more thoroughly than longer ones. As a first step, we used the maximum sequence length (1024) supported by some predictors (e.g., ESMFold), resulting in a maximum window length of 1021. We then tested progressively smaller window lengths down to a minimum of 3, yielding 171 different window sizes overall to evaluate model performance.

### Secondary structure assignment

As previously mentioned, the DSSP program by Kabsch and Sander [5] was utilized to assign secondary structure for all proteins to ensure consistent results in both predictions and experimental data.

The secondary structures assigned by DSSP are as follows.

- 3-turn helix (3 10 helix). A helix-like structure with a minimum length of 3 residues. This class is denoted by the letter G.
- 4-turn helix (*α* helix). A helix-like structure with a minimum length of 4 residues. This class is denoted by the letter H.
- 5-turn helix (*π* helix). A helix-like structure with a minimum length 5 residues. This class is denoted by the letter I.
- Hydrogen bonded turn (3, 4 or 5 residue turn). Forms a turn-like structure and is denoted by the letter T.
- Extended strand in a *β*-sheet conformation forming a pleated sheet structure. This class is denoted by the letter E.
- Residue in an isolated *β*-bridge (single pair *β*-sheet hydrogen bond formation). This class is denoted by the letter B.
- Bend (the only non-hydrogen-bond based assignment). This class is denoted by the letter S.
- Coil (none of the above), denoted by the letter C.

All recent experimental data is available in the macromolecular crystallographic information file (mmCIF) format since 2019 [50]. As a result, we utilize the modern format and a modernized version of DSSP program which include additional secondary structure classes and ability to handle recent experimental data formats. The differences in secondary structure classification in the modernized DSSP, e.g. the addition of polyproline helices, are ignored for Q8 prediction compatibility. Further discrepancies from the original version of DSSP are detailed in Section 2 of Supplementary materials.

Tertiary structure prediction methods typically output results in ‘PDB’ format, though some, like AlphaFold2, also support mmCIF format. For consistency, we used the ‘PDB’ output for all methods. To align with the process used for experimental structures, we converted the predictions to mmCIF format using MAXIT (https://sw-tools.rcsb.org/apps/MAXIT). We then applied DSSP to assign secondary structures to the mmCIF-formatted predictions.

### Secondary structure metrics

To assess the performance of secondary structure prediction methods on Q8, three commonly used metrics are employed: Accuracy (*Q*^*Acc*^) [11], Segment Overlap (*SOV*), and SOV_REFINE [14]. We refer to Segment Overlap as SOV99 [13], which improved the scoring from its original inception [12]. Its most recent modification is referred to as the refined version SOV_REFINE [14]. These metrics are computed using our “Secondary Structure Metrics Calculator” software (https://github.com/ivanpmartell/SSMetrics), which efficiently implements previously stated secondary structure metrics [11].

The metrics require two secondary structures of length *n* as input:

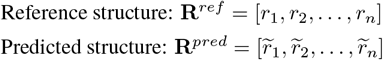

For the protein sequence **S** = [*a*_1_, …, *a*_*n*_] each amino acid *a*_*i*_ is assigned, either by DSSP or through prediction methods, a secondary structure *r*_*i*_ resulting in a secondary structure **R** = [*r*_1_, *r*_2_, …, *r*_*n*_]. Each structure contains a subset ϒ_*ref*_, ϒ_*pred*_ ϒ_8_ of the eight possible DSSP-assigned secondary structure classes ⊆ ϒ_8_ = {ℬ, 𝒞, ℰ, 𝒢, ℋ, ℐ, 𝒮, 𝒯}. For each SSE class *r*, a *reference segment* (or *reference r-block*) in **R**^*ref*^ is defined as a contiguous substructure 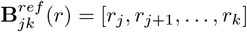 of **R**^*ref*^ that satisfies the following conditions:

Uniform Structure: All residues in the substructure are of SSE class *r* (i.e., *r*_*j*_ = *r*_*j*+1_ = … = *r*_*k*_ = *r*). Boundary Conditions: *r*_*j*−1_ ≠ *r* and *r*_*k*+1_ ≠ *r*, thus the SSEs immediately before *r*_*j*_ and after *r*_*k*_ must not be SSE class *r*.

Similarly, for each SSE class *r*, a *prediction r-block* in **R**^*pred*^ is defined as 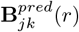 following the same criteria. The sets of all such blocks are defined as:

Reference Blocks:

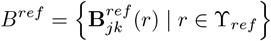

Prediction Blocks:

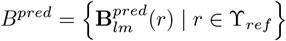

Here, ϒ_*ref*_ represents the set of all secondary structure element (SSE) classes present in the reference structure sequence **R**^*ref*^.

All secondary structure metrics evaluate SSEs at each position of both reference and predicted structure sequences using the *identity* function

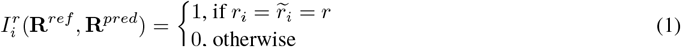

where *i* denotes the position of the SSE in both secondary structures **R**^*ref*^ and **R**^*pred*^, and *r* is the SSE class.

The Accuracy metric is defined as the ratio of matching SSE pairs between the reference and predicted structures to the total number of SSE pairs. Since the reference and predicted structures have equal lengths, the total number of SSE pairs is equal to the length of the structure sequences, i.e., |*R*^ref^| = |*R*^pred^| = *n*. This is more formally defined as:

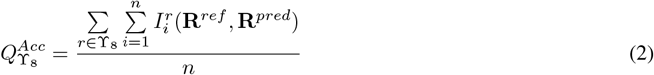

Accuracy, which measures exact matches between two secondary structures at each position, may not be able to sufficiently capture the structural details of slightly misaligned secondary structure elements (SSEs) that extend across the structure. To address this limitation, a more informative metric called Segment Overlap (*SOV*) was introduced [12] and consequently improved [13].

The Segment Overlap metric is a weighted sum over overlapping pairs of segment blocks for each SSE class *r* ∈ ϒ_*ref*_, accounting for slight misalignments in SSEs. Formally, for *r*-blocks 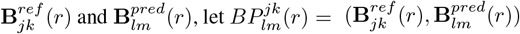 be an **overlapping segment block pair** for **R**^*ref*^ when *j* ≤ *m* and *l* ≤ *k*. Then, let 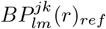 denote the first element of 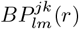 (i.e., 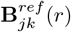)) for that overlapping segment block pair.

We denote 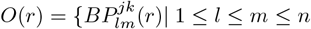 and 1 ≤ *j* ≤ *k* ≤ *n*} as the **set of all overlapping pairs** of *r*-blocks between **R**^*ref*^ and **R**^*pred*^ (i.e., the set of all tuples 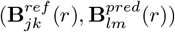 for all *r* ∈ ϒ_*ref*_). If a reference *r*-block 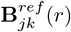 has no overlap with any predicted *r*-block in *B*^*pred*^, we define 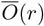 as the set of non-overlapping segment blocks in **R**^*ref*^ for SSE class *r*:

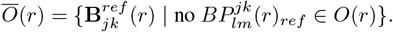

We now specify the functions that take part in the *SOV* definition.

Norm(*r*) is the **normalization value** for SSE class *r* defined as,

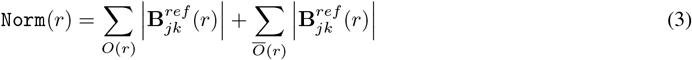

where 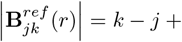 is the length of the *r*-block. It is important to note that any particular reference *r*-block can appear multiple times across different block pairs in *O*(*r*).

LenOv_*r*_ is the number of identical SSEs of class *r* for a pair of segments:

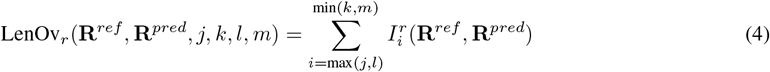

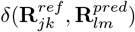 is the amount of **allowable misalignment** given to a pair of segments and defined as,

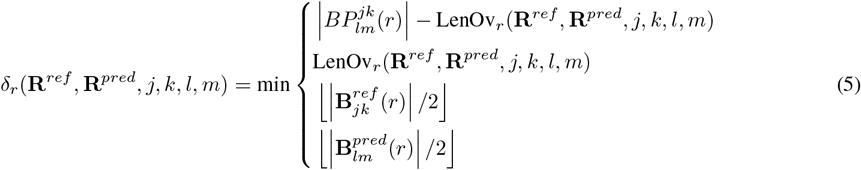

where 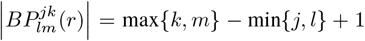 is the combined overlap length of an overlapping pair of segment blocks.

To define **Segment Overlap**, it is important to note that the set of all overlapping pairs of *r*-blocks *O*(*r*) contain *r*-blocks with their respective starting and ending indices (e.g.,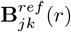, where *j* is the starting index of the *r*-block for **R**^*ref*^, and *k* is the ending index). Then for SSE class *r*, we define Segment Overlap *SOV* (*r*):

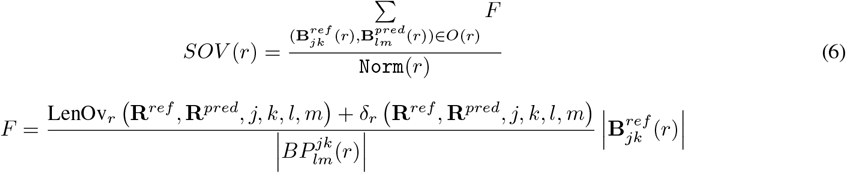

Lastly the overall *SOV* score for all SSE classes is defined as,

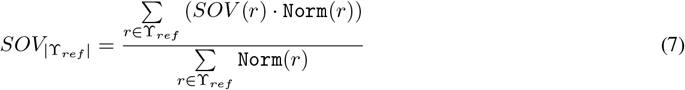

The refined version of *SOV*, termed as SOV_REFINE, changes the allowance function as follows,

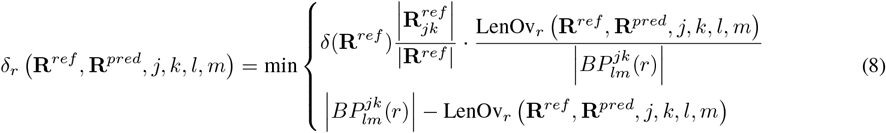

where δ (**R**^*ref*^) is the **total misalignment allowance** given for all segment blocks of the reference structure sequence as

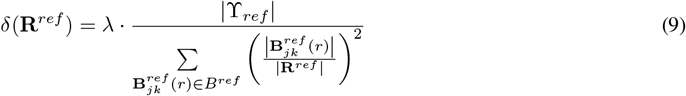

where ϒ_*ref*_ ⊆ ϒ_8_ and |ϒ_*ref*_ | is the number of SSE classes that appear in the reference structure sequence, and *λ* ∈ℝ, 0 ≤ *λ* ≤ 1 is an adjustable scale parameter that is used to limit the range of δ (**R**^*ref*^).

### Mutational metrics

Mutational data consists of a wild-type protein sequence along with its associated mutations. The previously discussed metrics are specifically designed to compare two secondary structures: a reference structure and a predicted structure. For each protein, the reference structure is derived using DSSP, while the predicted structure is generated by a structure prediction method.

To evaluate the performance of prediction methods on mutational data, it is essential to account for mutational changes. To achieve this, we utilize three mutational metrics, described below.

**Mutational consistency** measures the Accuracy between mutational changes in the reference and predicted structures. To compute this, we:

1. Compare the wild-type and mutated reference structures element-wise for each secondary structure element (SSE). This yields the **reference mutational change** sequence, indicating whether each SSE is preserved (‘N’) or changed (‘C’) after mutation.
2. Perform the same comparison for the wild-type and mutated predicted structures, producing the **predicted mutational change** sequence.
3. Calculate Accuracy for the reference and predicted mutational change sequences, treating the two possible states—’preserved’ and ‘changed’—as binary classes.

By framing mutational consistency through binary states, additional binary classification statistics [57] can be used to further analyze mutational data. These statistics can give further insights into mutational specific structural changes for the resulting positive (preserved) and negative (changed) classes.

**Mutational accuracy** measures Accuracy for SSE mutations in the reference and predicted structures. The calculation involves the following steps:

1. Compare the wild-type and mutated reference structures element-wise for each secondary structure element (SSE). This generates the **reference SSE mutation** sequence, indicating the type of change, e.g., ‘EE’ for no change in a *β*-strand or ‘EI’ for a *β*-strand changing into a *π*-helix.
2. Perform the same comparison for the wild-type and mutated predicted structures to produce the **predicted SSE mutation** sequence.
3. Calculate Accuracy for the reference and predicted sequences using the SSE change types from step 1 to replace the standard secondary structure classes.

For Q8, there are 64 possible SSE mutation classes, as each of the eight secondary structure classes can either remain the same or change into any other class after mutation.

**Mutational precision** is calculated as Accuracy, Segment Overlap, or SOV_REFINE between interlaced SSE sequences of the wild-type and predicted secondary structures. The process involves:

1. Interlacing the equal-length wild-type and mutated structures element-wise for each SSE. For a wild-type secondary structure **R**^*rep*^ = [*r*_1_, *r*_2_, …, *r*_*n*_] and a mutated structure 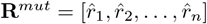, the **interlaced SSE sequence** is defined as:

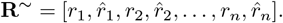
2. This procedure is applied to both the reference structure and predicted structure, yielding the **reference interlaced SSE** and **predicted interlaced SSE**.
3. Finally, Accuracy, Segment Overlap, or SOV_REFINE is computed between the reference and predicted interlaced SSE sequences.

To mitigate potential confusion, we provide an illustrative example, showing results for all the secondary structure metrics described above in Table 1 with the following data,

**Table 1:**
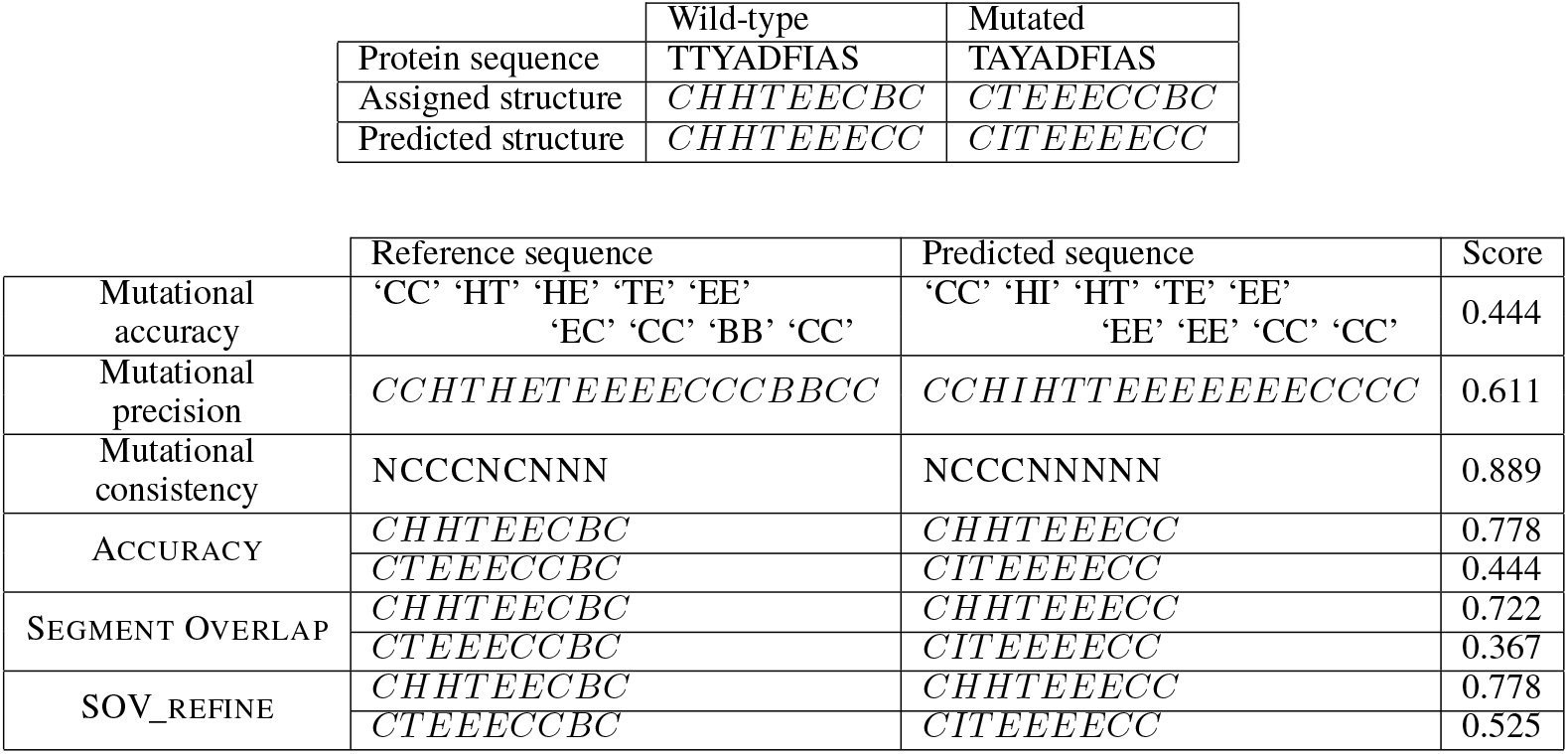
Metric results for example data.

### Prediction method spread

Prediction methods can also be assessed by the distribution spread of their performance along different secondary structure metrics. To compare different prediction methods, their evaluation must be measured on a common set of protein sequences **S**^∗^. For our purposes, this dataset should contain protein sequences for which a prediction method *m* will predict a secondary structure. The secondary structure assignment to these sequences is also required, and are obtained through DSSP.

Formally, let **A** = {*A, C, D, E, F, G, H, I, K, L, M, N, P, Q, R, S, T, V, W, Y*} be a finite set containing the alphabet of standard amino acids as defined by residue or side chain. A **protein sequence** over **A** is a finite sequence of amino acids **S** = [*a*_1_, …, *a*_*n*_] of length *n*, where *a*_*i*_ ∈ **A**, for all 1 *i* ≤ *n*. ≤*a*_1_ is the N-terminus residue and *a*_*n*_ is the C-terminus residue. For a protein sequence **S** each amino acid *a*_*i*_ is assigned to a secondary structure element *r*_*i*_ resulting in the complete protein secondary structure **R** = [*r*_1_, *r*_2_, …, *r*_*n*_]. Then, every secondary structure element must be assigned a DSSP class *r*_*i*_ ∈ ϒ_8_, where ϒ_8_ = {𝒞, ℋ, ℰ, 𝒢, ℐ, 𝒯, 𝒮, ℬ} is the set of possible SSE classes in DSSP.

Mutational metric results compared to non-mutational metrics. The top table shows the protein data given as an example to utilize in the computation of the metrics. The bottom table shows the input reference and predicted sequences used while computing the metric score. For mutational metrics, the input is transformed as detailed in the Mutational metrics section.

A prediction method *m* is defined as a learnable function that maps the protein sequence **S** to a predicted secondary structure sequence 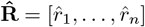, where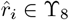 :

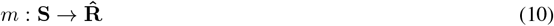

The function *m* can be represented as any machine learning model that takes a protein sequence **S** as input and outputs a predicted secondary structure sequence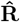.

Let the results dataset 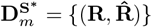 for prediction method *m* be a set of tuples containing assigned and predicted sequences for each protein sequence **S** ∈ **S**^∗^, for **S**^∗^ ={**S**_1_, …, **S**_*z*_}. Then, each tuple can be scored by any secondary structure metric defined above, where **R**^*ref*^ = **R** and 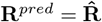.

The standard deviation is a conventional measure of spread for a distribution when its mean is the most appropriate measure of the distribution centre. Simply using standard deviation as a measure of distribution spread does not account for the performance of the prediction method. This can lead to low performing prediction methods with a low standard deviation to be considered superior to higher performing prediction methods. To account for this, we define two types of spread for a results dataset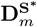. We start with Extreme spread, *XTSpread*, defined as,

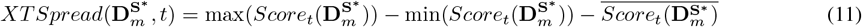

where the *Score*_*t*_ function returns a list of scores calculated from a secondary structure metric *t* for every tuple in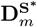. As the dataset *D* contains multiple proteins and a prediction for each protein, the secondary structure metric scores for all the predictions result in a distribution of scores over *D* for each prediction method. Thus, we can obtain the maximum value max(*Score*(*M, D*)) and minimum value min(*Score*(*M, D*)) for the distribution, along any distribution statistic such as its mean value, 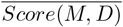, and standard deviation, *σ*(*Score*(*M, D*)). *XTSpread* is defined as the difference between the distribution’s range and its mean. This measure decreases when the spread of the data is narrower and the mean is higher. It captures the most extreme values—the best and worst predictions—making it particularly useful for assessing low-confidence predictions and quantifying the potential impact of large errors. Similarly, we define Standard spread, *STSpread*, as,

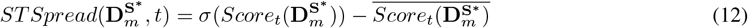

which, like the *XTSpread*, returns a lower value when the spread is narrower and the mean is higher. However, because *STSpread* does not account for the entire distribution range, it excludes extreme values. Consequently, *STSpread* captures the overall performance of a prediction method without considering outcomes distributed outside a single standard deviation of its mean.

## Results

This study initially examines each top-, average-, and low-performing predictor, grouped according to their significance in three different feature selection algorithms. In previous work [11], each predictor’s performance was determined based on its mutational capabilities, which confirmed the importance of each group. Specifically, the top-performing predictors include AlphaFold2, ColabFold, ESMFold, and SSPro8; the average-performing predictors include SPOT1D and SPOT1D-LM; and the low-performing predictors include SPOT1D-Single, RGN2, and Raptor-X Property.

In addition, we compare different machine learning algorithms that can be used during our refinement strategy. These comparisons justify our choice of algorithm for creating the proof-of-concept ensemble model, Mut2Dens. The creation of Mut2Dens is intertwined with our refinement strategy, and as such we describe the this strategy alongside Mut2Dens. We then show the performance results on test data using the best-performing ML algorithm and input representation combination that formed our ensemble model, along different versions of Mut2Dens depending on its input predictors. The multiple versions of Mut2Dens allow us to investigate how different types of predictors can affect the refinement output. Finally, we provide an in-depth analysis of the ensemble approach, illustrating how the ensemble model refines the secondary structures and its potential benefits and drawbacks.

### Feature selection

Feature selection algorithms are highly dependent on the data representation they receive. As with tree-type ML algorithms, the inputs can be structured in various ways. In our initial strategy, we used nominal data with the full sequence length for each prediction method. Each method’s feature importance was calculated by summing the contribution of all secondary structure elements (SSEs) it assigned, effectively weighting each method by its cumulative SSE impact. However, as shown in Fig 3 A, the algorithms had difficulty discerning feature significance under this approach, and no clear consensus emerged across different feature selection methods.

**Figure 3.**
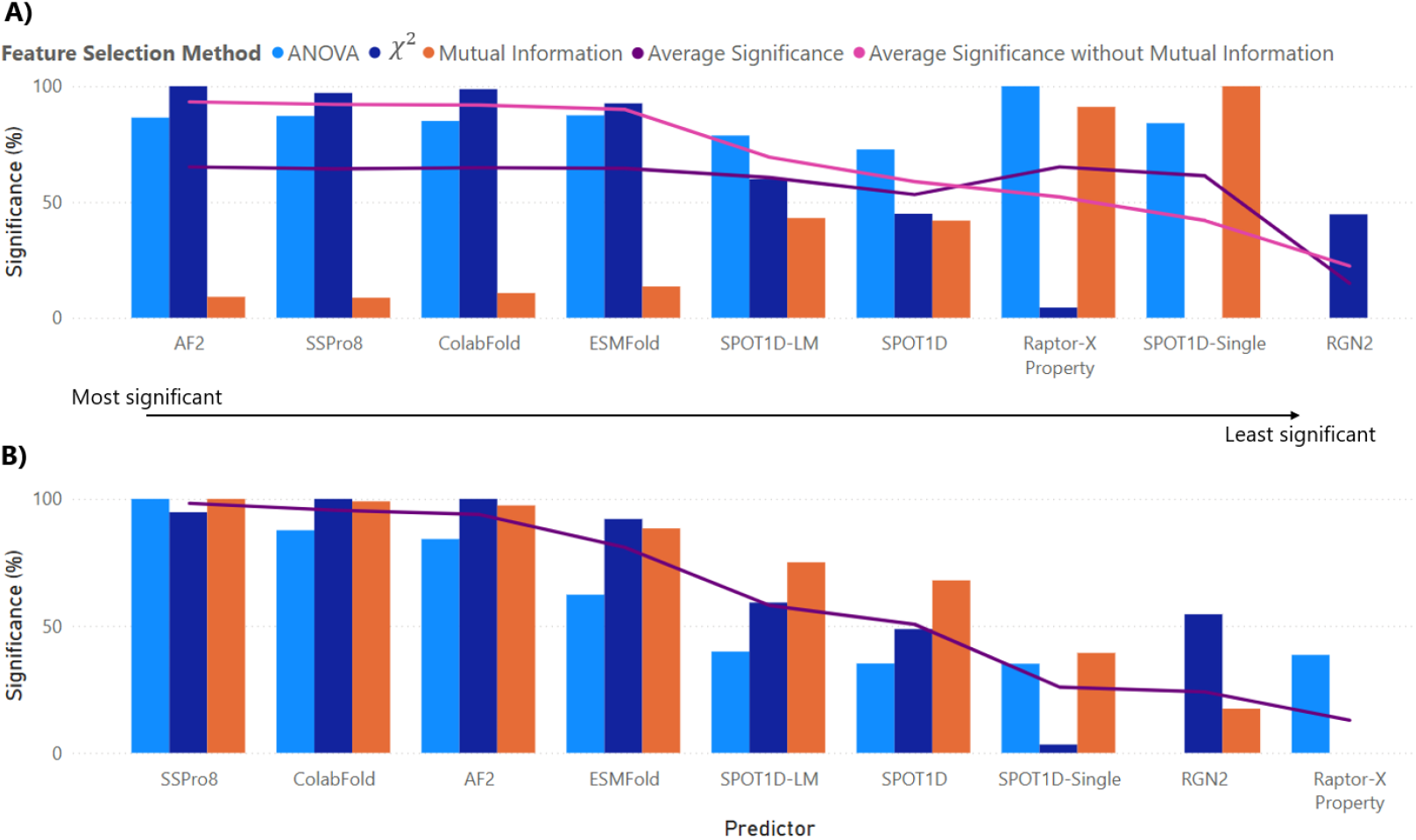
Feature selection. Score percentage for three differing feature selection algorithms: ANOVA, *χ*^2^, and Mutual Information. Both graphs show the most significant predictors from left to the least significant predictors on the right. A)Results from nominal data. The purple line shows the average significance for all three algorithms for a specific predictor. The pink line shows the average for ANOVA and *χ*^2^. Removing Mutual Information gives similar results for both windowed and full sequence nominal data where top, avg, and low performing methods follow the same trend to their significance. B) Results obtained from windowed nominal data. The line shows the average significance for all three algorithms for a specific predictor.

To address this issue, we adopted a windowed input approach. This yielded more cohesive results, with the top predictors—AlphaFold2, ColabFold, ESMFold, and SSPro8—being consistently identified by all three feature selection algorithms (Fig 3 B), a finding that aligns with our earlier analysis [11]. Meanwhile, Raptor-X Property ranked among the least significant methods in both *χ*^2^ and Mutual Information tests, yet it appeared highly significant in the full-sequence analysis for ANOVA and Mutual Information. Consequently, we still included Raptor-X Property in our ensemble model to see if adding a lower-performing prediction method with minimal feature importance could nevertheless improve performance.

The feature selection process identified several key predictors that significantly influence the ensemble model’s performance. Unsurprisingly, their significance generally aligns with their performance scores, making the top-performing predictors the most important. Meanwhile, Mutual Information-based feature selection diverged considerably when using windowed sequence lengths, and ANOVA also varied with changes in data representation. Interestingly, when Mutual Information was excluded from the nominal data approach, the results mirrored the windowed nominal data findings, with top-performing predictors emerging as the most significant and low-performing predictors as the least significant.

### Machine learning algorithms

We compared the performance of tree-type models and neural-type models in predicting secondary structure. Details of the tree-type models and neural-type models are given in Section 5 of Supplementary materials. Results can be seen in Fig. 4. The tree-type models, such as decision trees and random forests demonstrated superior performance with higher average SOV_REFINE scores and narrower confidence intervals, indicating more consistent predictions. Specifically, the tree-type models achieved an average SOV_REFINE score of 95%. In contrast, the neural-type models, including convolutional neural networks (CNNs) and recurrent neural networks (RNNs), showed lower average SOV_REFINE scores and wider confidence intervals, reflecting greater variability in their predictions. The best neural-type model (transformer) achieved a similar average SOV_REFINE score of 93% to tree-type models. While neural-type models have the potential for improvement through extensive hyper-parameter tuning and improved architectural designs, tree-type models outperform them in terms of both accuracy and consistency for this limited mutational dataset.

**Figure 4.**
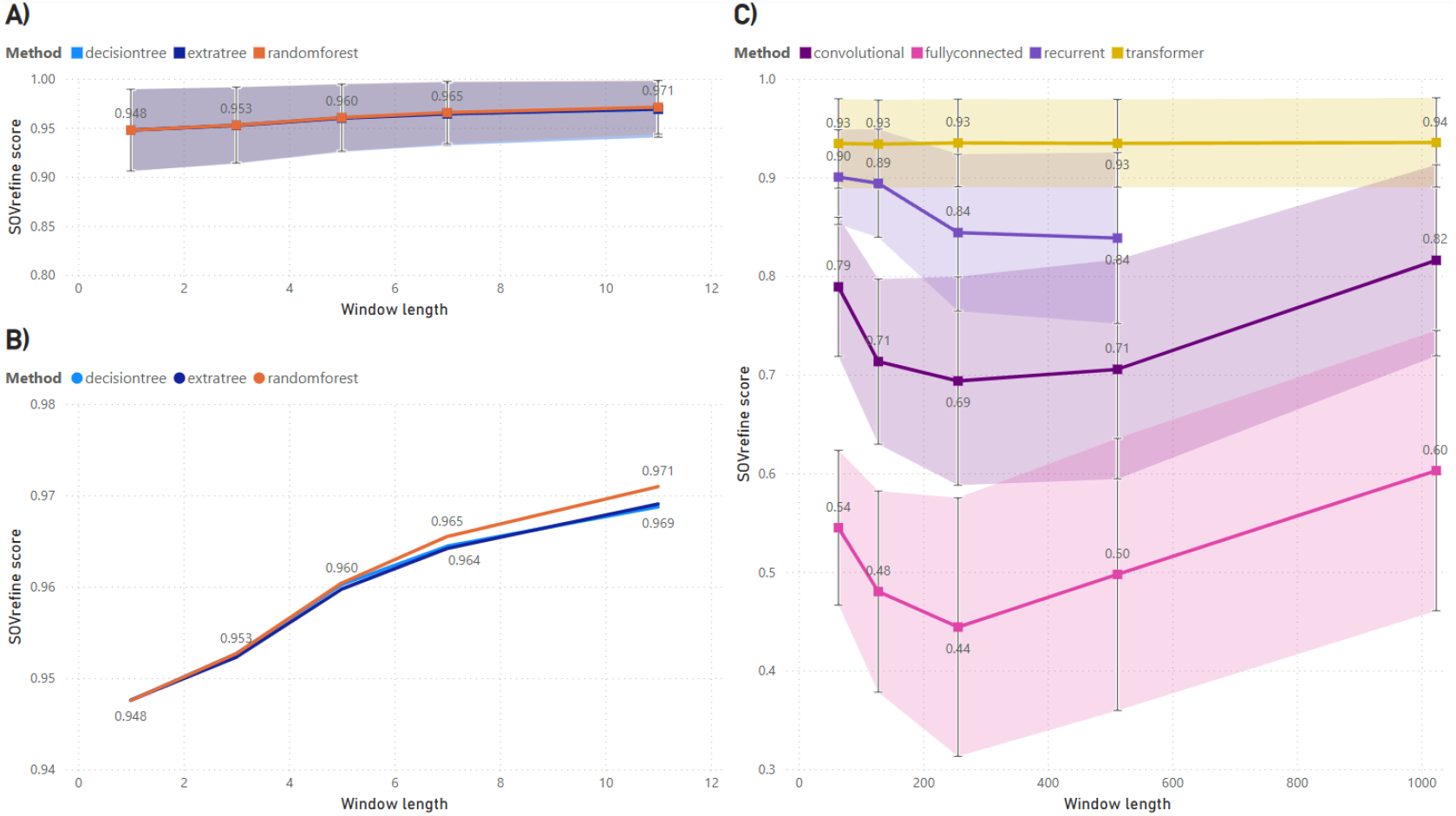
Comparison of Tree-type and Neural-type trained models. A) Results from tree-type models showing their average SOV_REFINE score and their confidence intervals using the 25^th^ and 75^th^ percentiles. B) A magnified look into the tree-type models for differing window lengths. C) Results from neural-type models, showing their average SOV_REFINE scores and confidence intervals with 33^rd^ and 66^th^ percentiles to reduce the interval overlap in the visual. Although tighter percentiles are used in neural-type models, confidence intervals are wider than tree-type models. Clearly, tree-type models outperform network-type models for this dataset. Further improvements to neural-type models should be possible but would require large amounts of hyper-parameter tuning and design considerations.

### Tree ensemble results

As previously demonstrated, tree-type models showed superior performance on our mutational dataset. We therefore focused on tree-type algorithms — specifically extremely randomized trees — because of their robustness to overfitting. To assess the reliability and generalizability of these models, we performed a 7-fold cross-validation on the mutational dataset using a leave-one-out approach; the number seven was chosen to ensure folds were of uniform size. The results from this cross-validation are shown in Fig. 5.

**Figure 5.**
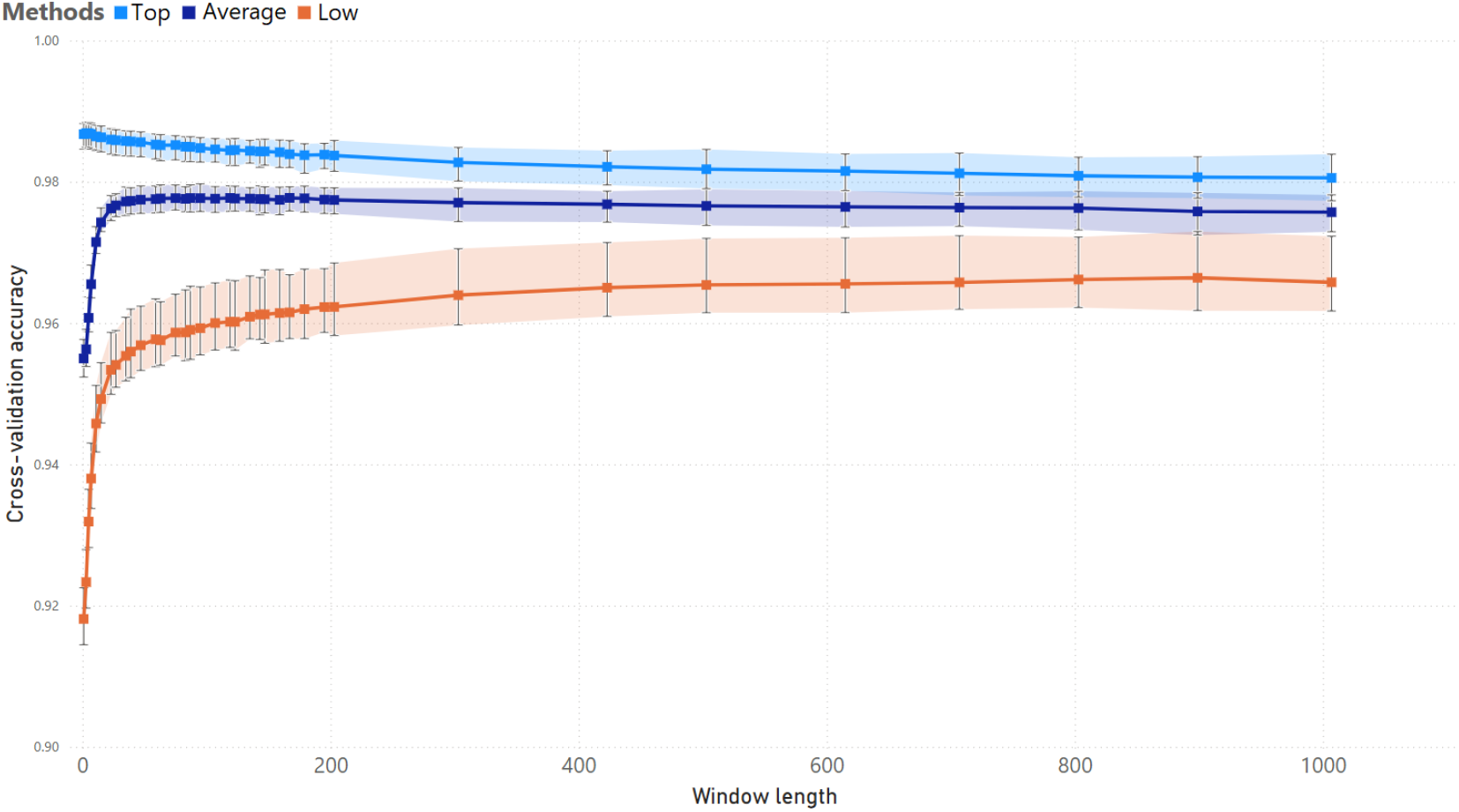
Cross-validation results. Further tree-type results using 7-fold cross validation with longer window lengths. The results are given for different input predictors: Top-performing, Average-performing, and Low-performing. Models created from Top-performing predictors show a slight decline in performance as window length increases, while the others improve as the window length increases.

We excluded tree-type algorithms that relied on a nominal data representation because they underperformed, possibly due to the discrepancy observed in the Mutual Information-based feature selection. It appears that absolute positional information imposed by nominal data does not yield meaningful insights into secondary structure. Consequently, all subsequent tree-type results presented here use a windowed nominal data representation.

An extremely randomized tree ensemble demonstrated consistent performance across all folds, achieving an average accuracy of 98% across all predictors when using longer window lengths. Interestingly, as predictors improve in performance and gain higher feature significance, shorter window lengths can sometimes help avoid misclassifications of secondary structures. It is worth noting that overall accuracy tends to be high because the mutational dataset consists of proteins differing by only a single amino acid.

### Mut2Dens

Here we describe the trained model Mut2Dens, which is built upon the knowledge gained during feature selection and model determination. As the feature selection algorithms suggest, we chose the most significant predictors, with an average significance percentage of 75 or higher, for agglomeration into Mut2Dens. We also added Raptor-X Property for its ambiguous importance during feature selection. Its low significance on the windowed representation can give insights into ensemble results from adding less significant predictors. We opt for ColabFold in place of AlphaFold2, as ColabFold is a computationally efficient version of AlphaFold2 while maintaining a high correlation between their predictions. Therefore, Mut2Dens is tailored to be computationally efficient with its mixed use of fast and accurate predictors. Further details of Mut2Dens are given in Section 3 of Supplementary materials. The final list of predictors are as follows,

1. SSPro8
2. ColabFold
3. ESMFold
4. Raptor-X Property

The predictor outputs are converted into nominal windowed data and concatenated into a single input vector. This input vector is passed to the ExtraTree model, which returns the refined output one amino acid at a time. Finally, the refined outputs are concatenated to form the complete refined secondary structure for the given sequence. A depiction of this process is shown in Fig 6.

**Figure 6.**
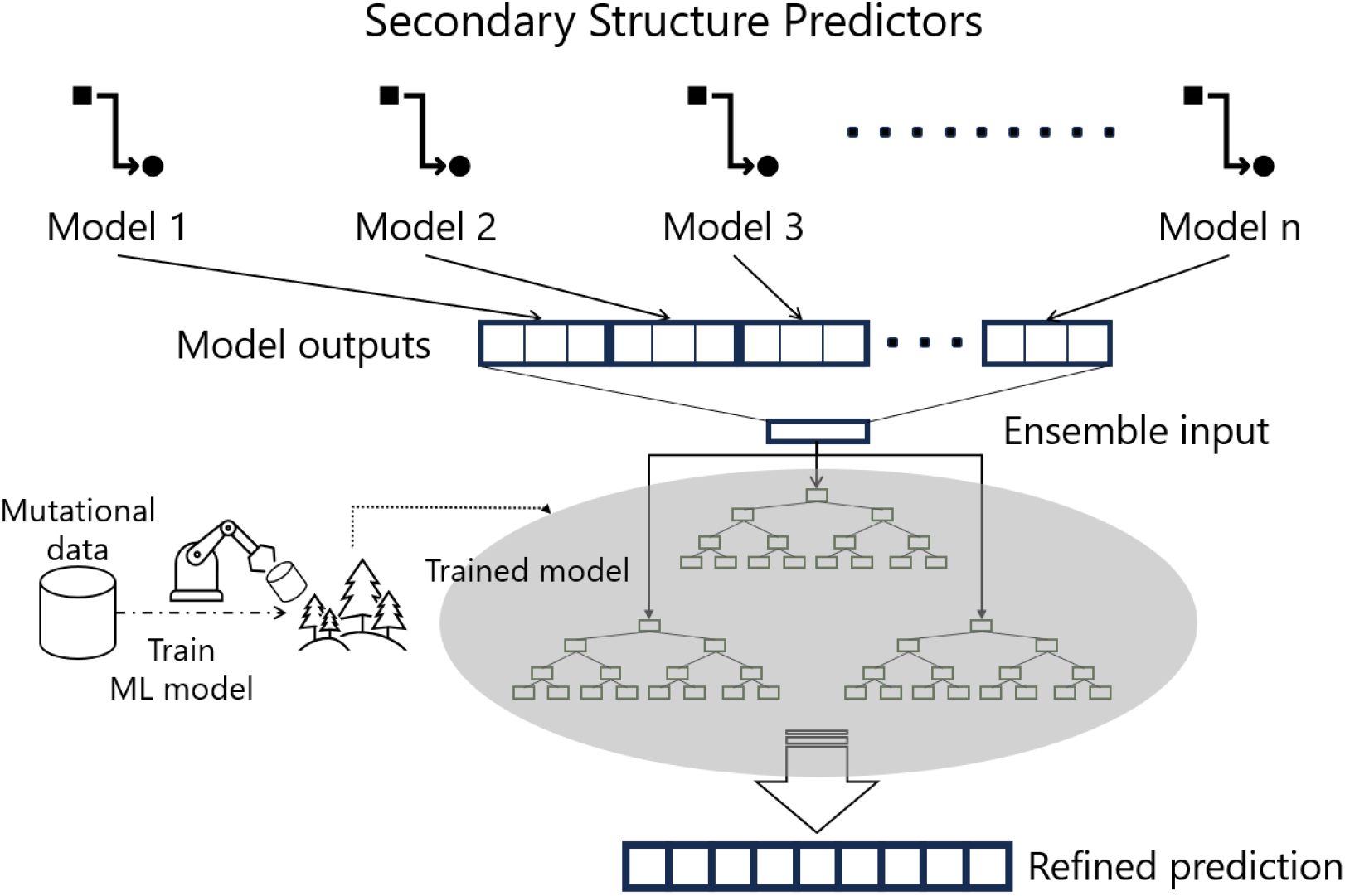
Refinement strategy. Diagram depicting the creation process of an ensemble model using our refinement strategy. First, the selected predictors are used to predict secondary structure for a given protein sequence. The predictors’ outputs are concatenated and used as input for a trained tree-ensemble model of extremely randomized trees. The trained model, Mut2Dens, outputs a refined prediction of the secondary structure, which takes into account its mutation-specific training.

### Mutational data results

Mutational capabilities of the models were evaluated using the mutational dataset along the mutational metrics described previously. The mutational precision and accuracy results, depicted in Fig. 7 A and B respectively, show that most models consistently achieved high scores across all metrics. These high results arise from the high degree of overlap between the mutational dataset and the training of these models. To minimize any potential overlap of the results and training data for Mut2Dens, results are shown for its testing dataset. While their mean results are high, the extreme values indicate a wide spread where low-confidence predicted proteins result in low predictive performance for most models. Our refinement strategy manages to reduce this spread, indicating the value of mutational refinement.

**Figure 7.**
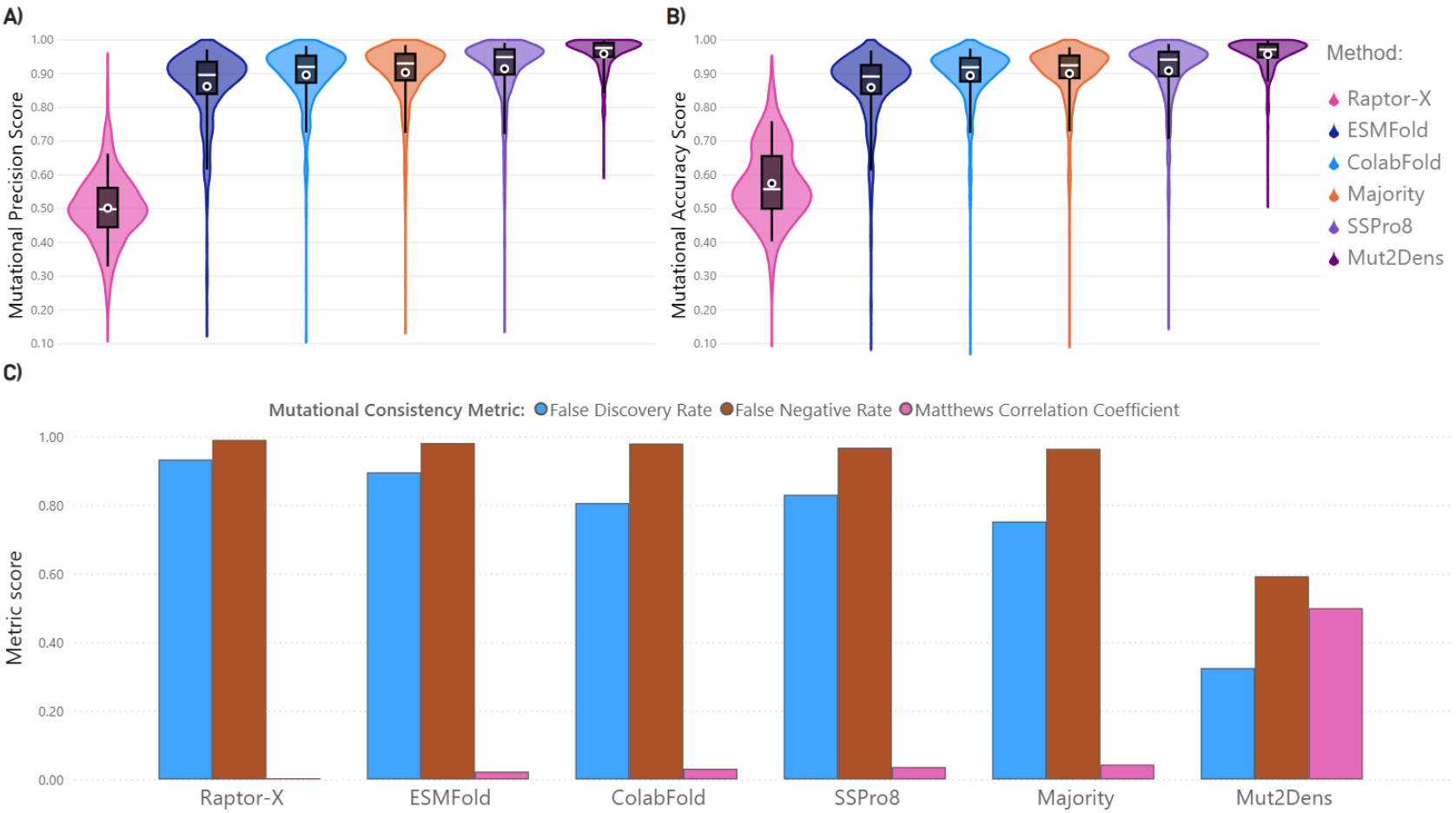
Mutational dataset results. Graphs showing mutational metric results for predictors, a majority agreement model, and Mut2Dens. Results show a narrower spread in the performance distribution, reducing the amount of highly incorrect predictions for Mut2Dens for the following metrics: A) Mutational precision, and B) Mutational accuracy. C) Results for mutational consistency metrics: False Discovery Rate (FDR), False Negative Rate (FNR), and Matthews Correlation Coefficient (MCC). Mutational consistency scores indicate whether the structural mutation occurs in the correct place. High values of FDR and FNR indicate poor performance in the model predicting the correct structural change location. MCC indicates the overall performance of the model, where higher is better

Mutational consistency converts the secondary structure classification problem into a two-class mutational classification task. For each amino acid location in the protein sequence, we ask whether the mutation caused a structural change or the structure remained stable. Fig. 7 C shows a high false negative rate for all predictors, including its majority consensus model. The false negative rate in this context denotes an incorrect prediction of a change when no such structural change occurs. Likewise, the high false discovery rates indicate that out of all structural changes predicted, most structural changes did not actually occur. Mut2Dens produces lower false discovery rates and false negative rates than the predictors, demonstrating the usefulness of mutational refinement. The low Matthews Correlation coefficient scores indicate low predictive performance of the model regarding the mutational classification task. We can clearly see that our refinement strategy can enhance the overall mutational predictive capabilities of predictors, in contrast to a simple combination like a majority agreement of predictors or the predictors themselves.

### Non-mutational data benchmarks

Mut2Dens was evaluated and compared to secondary structure predictors and a majority agreement model of the predictors. We also include an ablation study of the selected predictors for Mut2Dens that contained the best predictors for the test datasets, ColabFold and SSPro8. Therefore, multiple versions of Mut2Dens with different input predictors are shown below.

Performance of the Mut2Dens model was evaluated using the CB513 dataset, a widely recognized benchmark for secondary structure prediction for its non-homologous proteins, with a sequence similarity of less than 25% between all proteins. The results, depicted in Fig. 8, demonstrate that Mut2Dens achieved high SOV_REFINE scores, indicating its ability to predict secondary structures accurately. Specifically, the model outperformed all other predictors, including SSPro8, in terms of extreme value predictions, as shown by the XTSpread metric. When looking at standard (near the mean) protein predictions, Mut2Dens exhibited a non-significant decrease of performance of less than 1% compared to SSPro8, as indicated by the STSpread metric. Overall, the CB513 test benchmark results highlight the robustness and effectiveness of Mut2Dens in handling a diverse set of proteins. The capabilities of Mut2Dens are comparable to the best predictor for the dataset, while increasing the performance of the most inaccurate predictions for other predictors.

**Figure 8.**
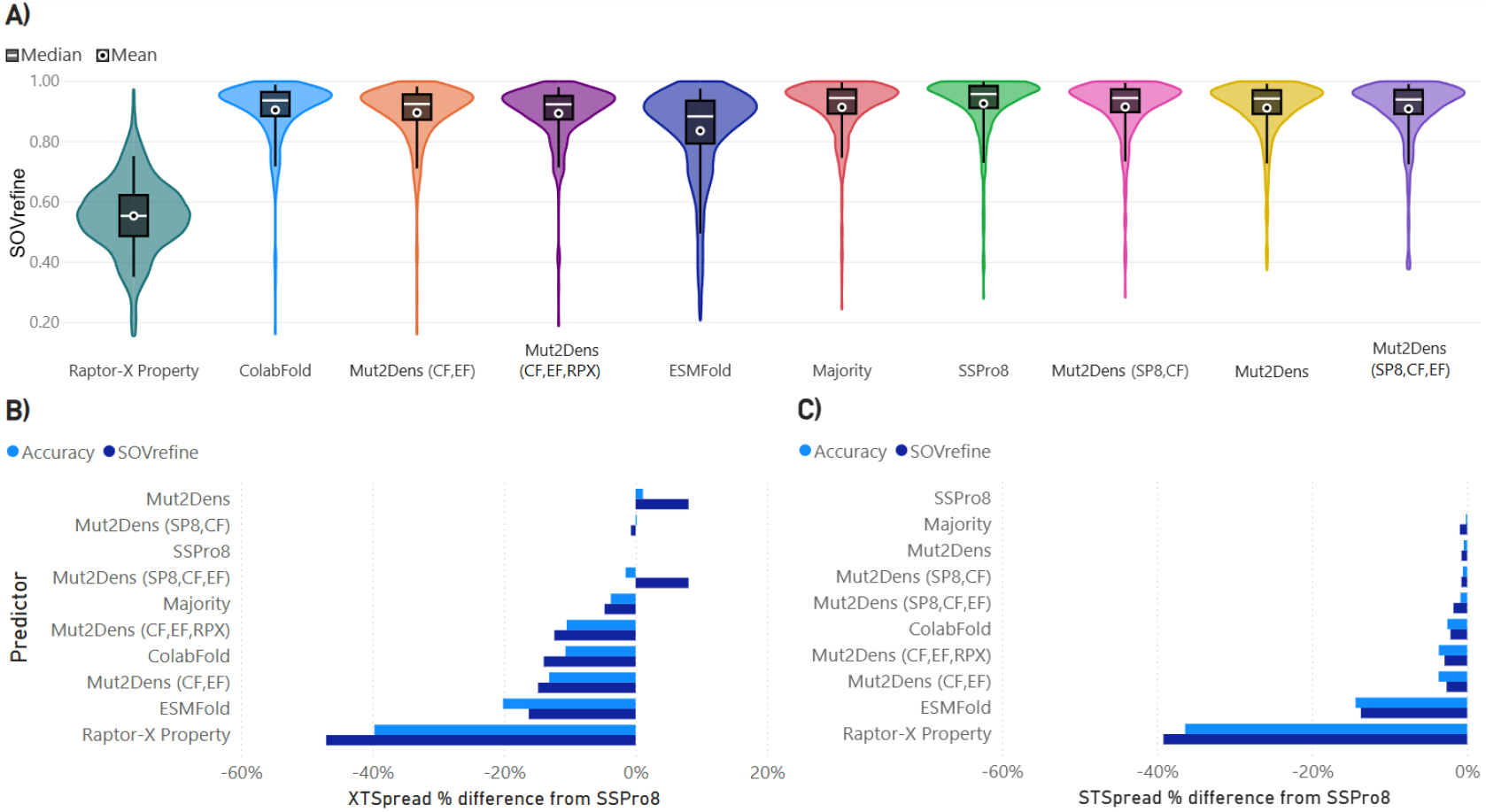
Testing models on CB513. This dataset has been previously utilized as a testing benchmark for many studies. Predictors utilized for ensemble models include ColabFold (CF), SSPro8 (SP8), ESMFold (EF), and Raptor-X Property (RPX). A) SOV_REFINE score results. The high scores result from most models utilizing this dataset. B) XTSpread difference to the best performing non-ensemble predictor for this dataset, SSPro8. C) STSpread difference to SSPro8. Taking the extreme values into account with XTSpread, we can see our ensemble model is capable of outperforming all others. Conversely, our ensemble model has a slightly lower performance than SSPro8 when focusing on non-extreme (very low performing) proteins.

From the different versions of Mut2Dens, we can see that the addition of predictors with low significance do increase the outcome slightly. It is likely that additional inclusion of uncorrelated predictors with differing methodologies will increase the outcome, although marginally. Therefore, the inclusion of such predictors will depend on a cost-effectiveness evaluation as each additional predictor increases processing time in a non-trivial manner.

The performance of Mut2Dens was also evaluated using the CASP15 dataset. The results for this benchmark dataset are shown in Fig. 9. Mut2Dens predictive capabilities increased the extreme values by a wide margin by almost 30%. This increase in low-performing predicted proteins come at a slight cost where the STSpread accuracy score of Mut2Dens decreases by about 5%. Most inaccurate predictions by secondary structure predictors have low confidence values. Mut2Dens is suggested to be utilized for these instances as its predictions can increase the structural outcome for such proteins and simultaneously provide insights from multiple predictors.

**Figure 9.**
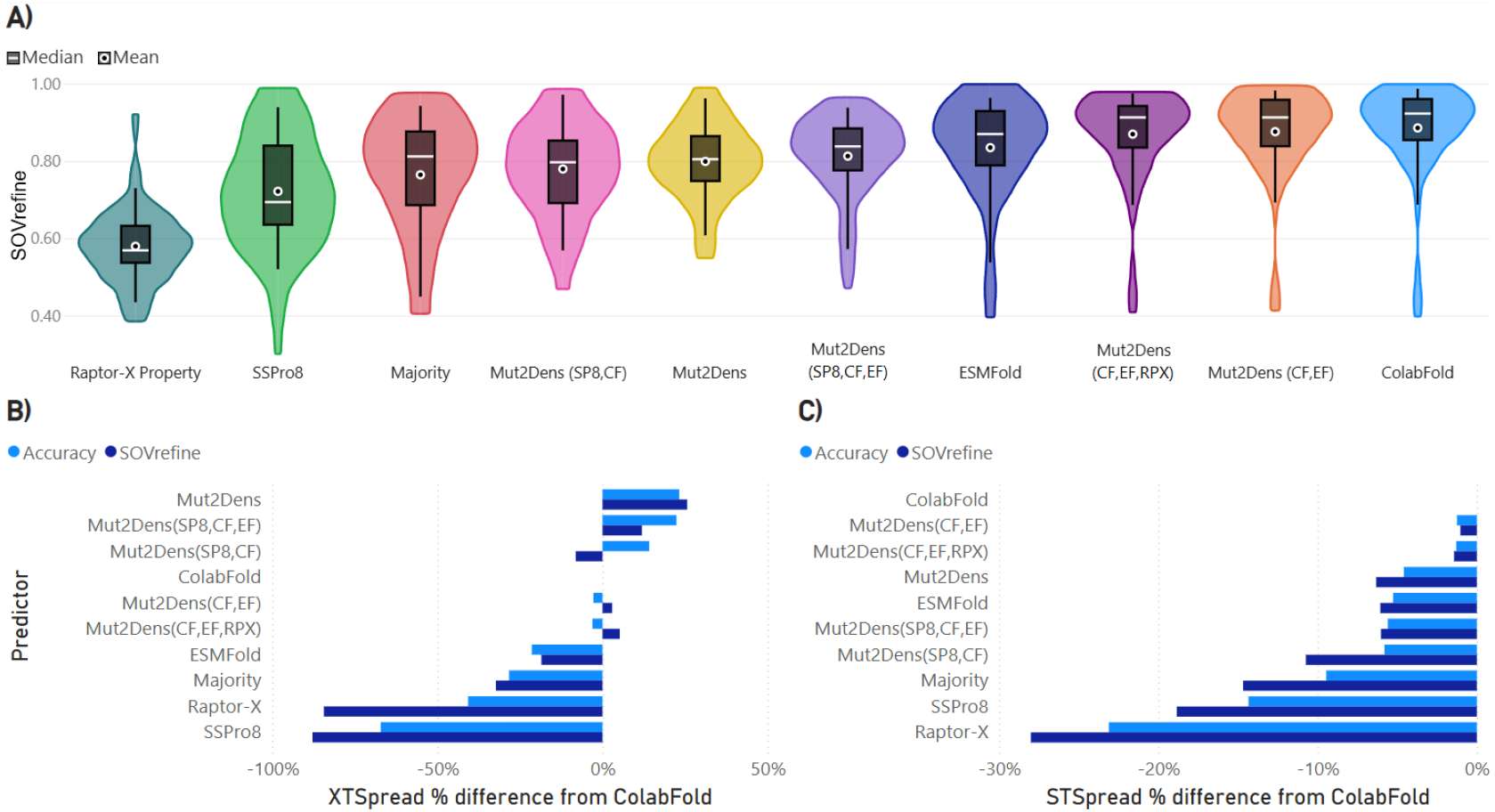
Testing models on CASP15. Most recent dataset with proteins that have not been included in the training of any model. Predictors utilized for ensemble models include ColabFold (CF), SSPro8 (SP8), ESMFold (EF), and Raptor-X Property (RPX). A) Performance of the models is more realistic than CB513 with a maximum mean SOV_REFINE of 88% by ColabFold. B) XTSpread and C) STSpread difference to the best performing non-ensemble model for this dataset, ColabFold. Similarly to CB513, the ensemble models outperform others when extreme values are taken into consideration, but perform slightly lower for non-extreme values.

### Knowledge-based model

Examining how tree-based models use our mutational data to refine secondary structure shows that their decisions align with findings from our previous mutational study [11], indicating that single amino acid mutations rarely converthelices into sheets (or vice versa). This insight is drawn primarily from crystallographic data, which necessitates that proteins form ordered crystal structures. Consequently, the data may be incomplete, yet the frequency of secondary structural interchanges observed in predictions remains higher than what current experimental evidence suggests.

Through the training procedure of tree-type models, we can compute a confusion matrix of the resulting SSE classes separated by every final decision (leaf) within the tree. This allow us to see the amount of weight any SSE class has when the model is expecting to predict a certain SSE. Our tree-type models contain over 1000 such prediction decisions and as such we aggregate them to obtain a complete picture of the weight each SSE class gives for an expected outcome. Therefore, we obtained decisions from 10 tree-type models to obtain a generalized average weight for each SSE class during an expected decision outcome. We use a simplified ensemble model with a window length of 1 to isolate the decisions taken for each model at a specific amino acid. Analyzing these weights, we obtain the results shown in Table 2. This square matrix contain possible secondary structure classes, where rows contain the expected or ‘true’ SSE and columns contain the SSE classes decided by the tree-type model. These values can be normalized to obtain the probability of the tree deciding a specific SSE class for each of the assigned SSE classes. The diagonal represents the secondary classes that are correctly decided, while the non-diagonal values represent overlapping decisions with other classes. We can see that *π*-helices have the most confounding with other classes as it is selected 84.4% when it is expected. Interestingly, *π*-helices are never confounded with *β*-sheets, isolated *β*-bridges, or coils. The model is able to predict a *π*-helix for an amino acid only when it is certain that *β*-sheets, isolated *β*-bridges, or coils are not feasible for that amino acid. This follows the knowledge-based rule that *π*-helix never turn into *β*-sheets, bridges, 3 − 10 helices, or coils within our mutational data.

**Table 2:**
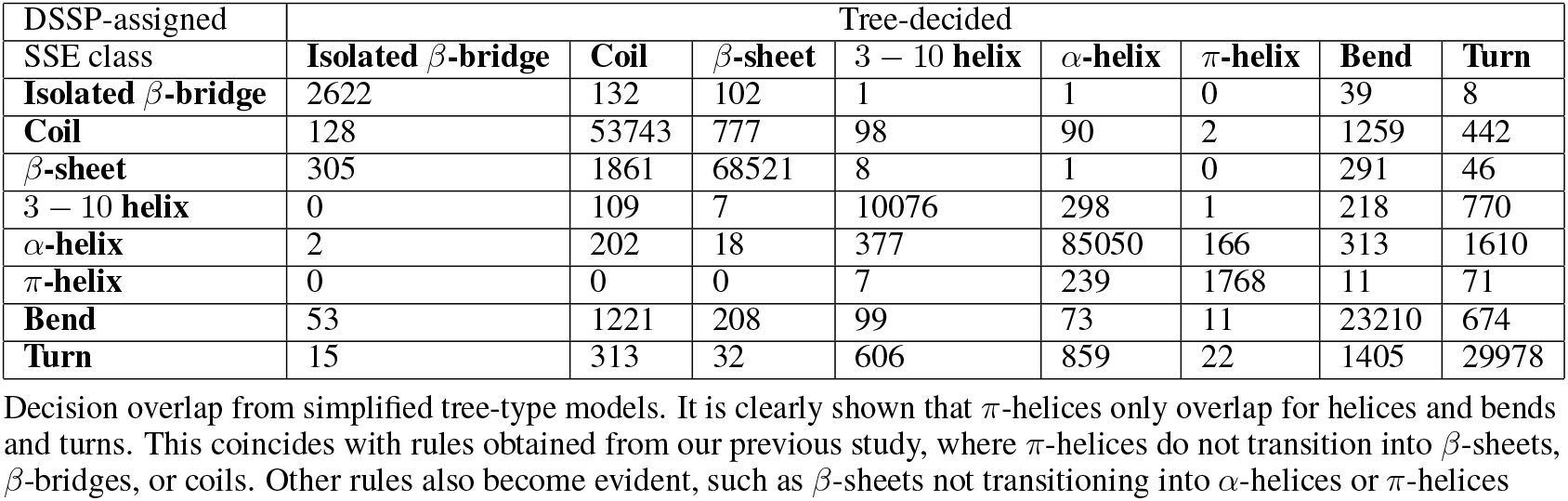
Tree knowledge.

Likewise, we can obtain all decision thresholds within the tree created during their training process. Aggregating these decisions thresholds allows us to obtain meaning from the decisions that the tree uses to obtain a certain outcome. The resulting human-readable rules, which simplifies the complexities of the tree-like structure, resemble the following example wording: “When ColabFold and ESMFold predict an *α*-helix, and SSpro8 predicts a *π*-helix, the ensemble outcome is *π*-helix.”

The advantage of using an ensemble model is the inclusion of multiple predictor outputs. Having different predictions allow us to compare and check which predictors have generated incorrect SSEs. We can also take these rules to see where the model generally thinks certain predictors require an adjustment. Furthermore, each individual protein prediction can also be compared to each predictor’s outcome for a more detailed view of the protein. Utilizing visualization tools such as 2dss (http://genome.lcqb.upmc.fr/2dss/), although limited to Q3 visuals, we can more easily see potential weaknesses of each predictor and have a better understanding of the resulting prediction from our ensemble model. An example of such a visualization is given in Fig. 10 comparing the DSSP-assigned structure, and predicted structures from Mut2Dens and the highly accurate ColabFold predictor. The numbers in the figure represent the location in the amino acid sequence, while the colors for the query describe the properties of an amino acid. The amino acid properties along with their color are the following,

**Figure 10.**
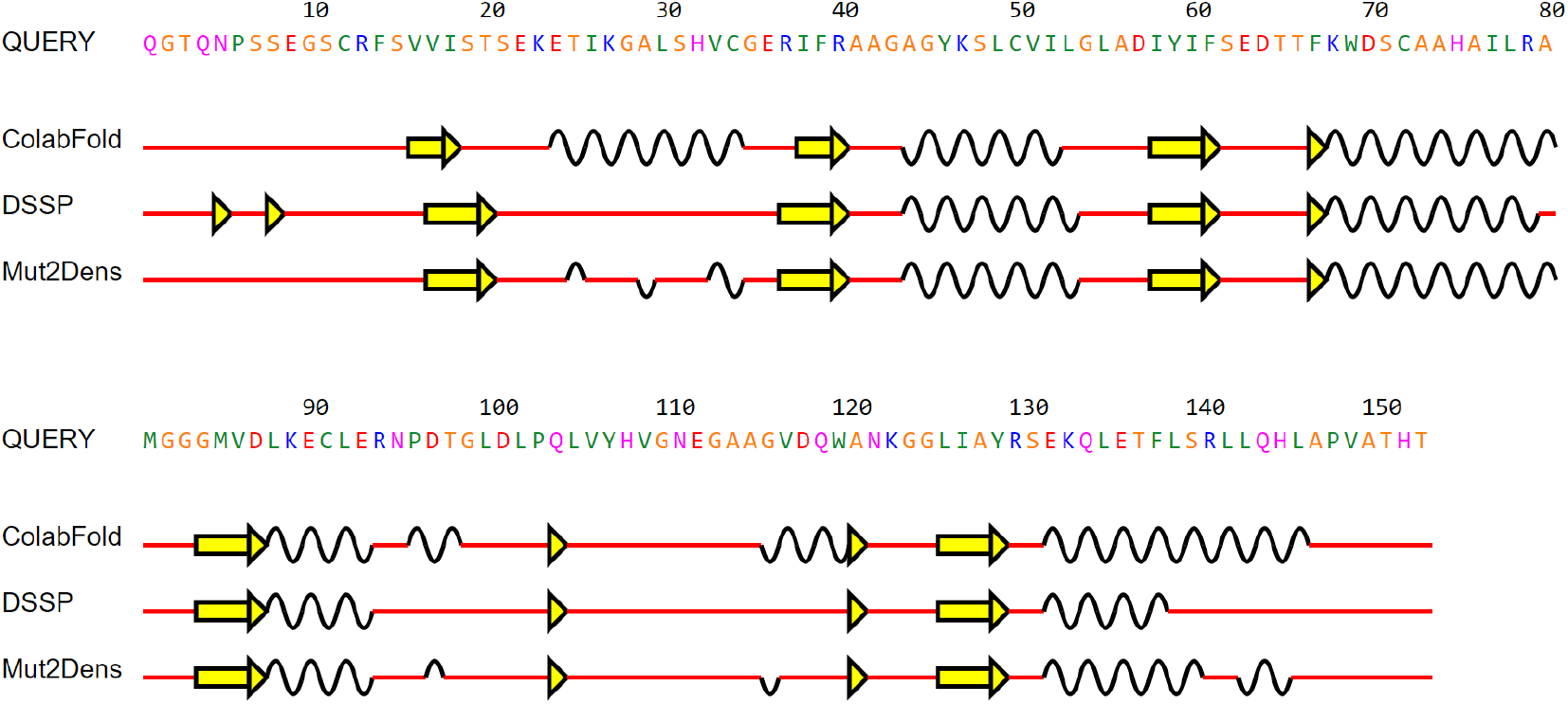
Structure refinement comparison. Visualization of predicted and assigned secondary structures. The secondary structure is simplified into Q3 for visualization purposes. For each amino acid, a line represents a coil, the yellow arrow represents a *β*-sheet, and the wavy line represents an *α*-helix. For this protein, ColabFold achieves 61% accuracy, while Mut2Dens achieves 82%. ColabFold predicts *α*-helices in several places that do not occur in the actual assigned structure by DSSP. While not perfect, Mut2Dens tries to correct these structures by removing most of the helical SSEs that are not part of the actual structure.

- Hydrophobic: green
- Small: orange
- Polar: magenta
- Negatively charged: red
- Positively charged: blue

## Conclusion

Although recent advances have led to highly accurate structure prediction methods, incorrect or low-confidence predictions are still possible — particularly when homology data are scarce. In such cases, these models may yield unreliable topologies. Furthermore, previous findings [11] show that current prediction methods often fail to predict the structural effects of single amino acid mutations accurately, sometimes producing highly improbable outcomes compared to experimental data.

To address these limitations, we developed a novel refinement strategy that relies solely on missense mutational data, requiring no additional information. By creating an ensemble of methodologically diverse predictors, we mitigate individual weaknesses while achieving reliable and consistent mutation-focused predictions. Our approach, implemented in the Mut2Dens model, integrates the strengths of selected predictors, thereby improving performance on mutational datasets without compromising results on non-mutational datasets.

Moreover, our refinement strategy enables the exploration or validation of biological insights through interpretable machine learning algorithms, such as tree-based models. Future work could extend these findings to tertiary structure predictions by leveraging knowledge gained from inconsistencies observed in secondary structure predictions. Additionally, we include the use of visualization tools that highlight discrepancies among predictors, revealing which secondary structures have been modified during refinement—thereby guiding researchers to regions where the predicted protein structure may be unstable or unreliable.

It is important to note that this refinement strategy is not intended to replace current prediction models. Instead, it serves as an auxiliary tool to verify and enhance the integrity of structure predictions, especially when predictor confidence is low. Incorporating mutational data not only refines existing models but also improves low-scoring predictions for non-mutational data, addressing a significant shortcoming of current prediction methods. For instance, in cases where ColabFold produces low-performing predictions, Mut2Dens often yields more reliable secondary structure representations. The ability to improve low-confidence predictions also allows meaningful comparisons across various structure prediction methods.

Ultimately, our work narrows the gap in low-scoring protein structure predictions, offering a more accurate and reliable framework that benefits biomedical research and drug design. By enhancing the consistency of protein structure predictions, we provide researchers with a comprehensive toolkit for making informed decisions about predicted structures. Moreover, secondary structure-focused applications, such as protein functional prediction methods [58], can directly benefit from these more reliable secondary structure predictions.

## Acknowledgments

We would like to express our sincere gratitude to Microsoft and University of Alberta for providing the computational resources required for this project. We are also grateful to have a grant from Natural Sciences and Engineering Research Council of Canada. Microsoft AI for Health supported us through their Azure grant, while University of Alberta allowed us to use their ‘Industry Sandbox and AI Computing’ entrepreneurship resources. Their generous contribution enabled us to perform large-scale predictions and analyses that were essential to our research.

## References

[1] Brian Kuhlman and Philip Bradley. Advances in protein structure prediction and design. Nature Reviews Molecular Cell Biology, 20(11):681–697, November 2019. ISSN 1471-0080. doi:10.1038/s41580-019-0163-x. URL https://www.nature.com/articles/s41580-019-0163-x. Publisher: Nature Publishing Group.

[2] John Jumper, Richard Evans, Alexander Pritzel, Tim Green, Michael Figurnov, Olaf Ronneberger, Kathryn Tunyasuvunakool, Russ Bates, Augustin Žídek, Anna Potapenko, Alex Bridgland, Clemens Meyer, Simon A. A. Kohl, Andrew J. Ballard, Andrew Cowie, Bernardino Romera-Paredes, Stanislav Nikolov, Rishub Jain, Jonas Adler, Trevor Back, Stig Petersen, David Reiman, Ellen Clancy, Michal Zielinski, Martin Steinegger, Michalina Pacholska, Tamas Berghammer, Sebastian Bodenstein, David Silver, Oriol Vinyals, Andrew W. Senior, Koray Kavukcuoglu, Pushmeet Kohli, and Demis Hassabis. Highly accurate protein structure prediction with AlphaFold. Nature, 596(7873):583–589, August 2021. ISSN 1476-4687. doi:10.1038/s41586-021-03819-2. URL https://www.nature.com/articles/s41586-021-03819-2. Number: 7873 Publisher: Nature Publishing Group.

[3] John M. McBride, Konstantin Polev, Amirbek Abdirasulov, Vladimir Reinharz, Bartosz A. Grzybowski, and Tsvi Tlusty. AlphaFold2 Can Predict Single-Mutation Effects. Physical Review Letters, 131(21):218401, November 2023. doi:10.1103/PhysRevLett.131.218401. URL https://link.aps.org/doi/10.1103/PhysRevLett.131.218401. Publisher: American Physical Society.

[4] Hilal Keskin Karakoyun, Sirin K. Yüksel, Ilayda Amanoglu, Lara Naserikhojasteh, Ahmet Yesilyurt, Cengiz Yakıcıer, Emel Timuçin, and Cemaliye B. Akyerli. Evaluation of AlphaFold structure-based protein stability prediction on missense variations in cancer. Frontiers in Genetics, 14, 2023. ISSN 1664-8021. URL https://www.frontiersin.org/articles/10.3389/fgene.2023.1052383.

[5] Wolfgang Kabsch and Christian Sander. Dictionary of protein secondary structure: Pattern recognition of hydrogen-bonded and geometrical features. Biopolymers, 22(12):2577–2637, 1983. ISSN 1097-0282. doi:10.1002/bip.360221211. URL https://onlinelibrary.wiley.com/doi/abs/10.1002/bip.360221211. _eprint: https://onlinelibrary.wiley.com/doi/pdf/10.1002/bip.360221211.

[6] Christophe N. Magnan and Pierre Baldi. SSpro/ACCpro 5: almost perfect prediction of protein secondary structure and relative solvent accessibility using profiles, machine learning and structural similarity. Bioinformatics, 30(18):2592–2597, September 2014. ISSN 1367-4803. doi:10.1093/bioinformatics/btu352. URL 10.1093/bioinformatics/btu352.

[7] Yuedong Yang, Jianzhao Gao, Jihua Wang, Rhys Heffernan, Jack Hanson, Kuldip Paliwal, and Yaoqi Zhou. Sixty-five years of the long march in protein secondary structure prediction: the final stretch? Briefings in Bioinformatics, 19(3):482–494, May 2018. ISSN 1477-4054. doi:10.1093/bib/bbw129. URL 10.1093/bib/bbw129.

[8] Tomer Sidi and Chen Keasar. Redundancy-weighting the PDB for detailed secondary structure prediction using deep-learning models. Bioinformatics, 36(12):3733–3738, June 2020. ISSN 1367-4803. doi:10.1093/bioinformatics/btaa196. URL 10.1093/bioinformatics/btaa196.

[9] Jisna Vellara Antony, Prayagh Madhu, Jayaraj Pottekkattuvalappil Balakrishnan, and Hemant Yadav. Assigning secondary structure in proteins using AI. Journal of Molecular Modeling, 27(9):252, August 2021. ISSN 0948-5023. doi:10.1007/s00894-021-04825-x. URL 10.1007/s00894-021-04825-x.

[10] Yuan Zhang and Celeste Sagui. Secondary structure assignment for conformationally irregular peptides: Comparison between DSSP, STRIDE and KAKSI. Journal of Molecular Graphics and Modelling, 55:72–84, February 2015. ISSN 1093-3263. doi:10.1016/j.jmgm.2014.10.005. URL https://www.sciencedirect.com/science/article/pii/S1093326314001648.

[11] Ivan Perez, Ulrike Stege, and Hosna Jabbari. Missense mutations: Backbone structure positional effects, December 2024. URL https://www.biorxiv.org/content/10.1101/2024.12.23.630208v1. Pages: 2024.12.23.630208 Section: New Results.

[12] Burkhard Rost, Chris Sander, and Reinhard Schneider. Redefining the goals of protein secondary structure prediction. Journal of Molecular Biology, 235(1):13–26, January 1994. ISSN 0022-2836. doi:10.1016/S0022-2836(05)80007-5. URL https://www.sciencedirect.com/science/article/pii/S0022283605800075.

[13] Adam Zemla, Ceslovas Venclovas, Krzysztof Fidelis, and Burkhard Rost. A modified definition of Sov, a segment-based measure for protein secondary structure prediction assessment. Proteins: Structure, Function, and Bioinformatics, 34(2):220–223, 1999. ISSN 1097-0134. doi:10.1002/(SICI)1097-0134(19990201)34:2<220::AID-PROT7>3.0.CO;2-K. URL https://onlinelibrary.wiley.com/doi/abs/10.1002/%28SICI%291097-0134%2819990201%2934%3A2%3C220%3A%3AAID-PROT7%3E3.0.CO%3B2-K. _eprint: https://onlinelibrary.wiley.com/doi/pdf/10.1002/%28SICI%291097-0134%2819990201%2934%3A2%3C220%3A%3AAID-PROT7%3E3.0.CO%3B2-K.

[14] Tong Liu and Zheng Wang. SOV_refine: A further refined definition of segment overlap score and its significance for protein structure similarity. Source Code for Biology and Medicine, 13(1):1, April 2018. ISSN 1751-0473. doi:10.1186/s13029-018-0068-7. URL 10.1186/s13029-018-0068-7.

[15] Chia-Tzu Ho, Yu-Wei Huang, Teng-Ruei Chen, Chia-Hua Lo, and Wei-Cheng Lo. Discovering the Ultimate Limits of Protein Secondary Structure Prediction. Biomolecules, 11(11):1627, November 2021. ISSN 2218-273X. doi:10.3390/biom11111627. URL https://www.mdpi.com/2218-273X/11/11/1627. Number: 11 Publisher: Multidisciplinary Digital Publishing Institute.

[16] Sheng Wang, Wei Li, Shiwang Liu, and Jinbo Xu. RaptorX-Property: a web server for protein structure property prediction. Nucleic Acids Research, 44(W1):W430–W435, July 2016. ISSN 0305-1048. doi:10.1093/nar/gkw306. URL 10.1093/nar/gkw306.

[17] Jack Hanson, Kuldip Paliwal, Thomas Litfin, Yuedong Yang, and Yaoqi Zhou. Improving prediction of protein secondary structure, backbone angles, solvent accessibility and contact numbers by using predicted contact maps and an ensemble of recurrent and residual convolutional neural networks. Bioinformatics, 35(14):2403–2410, July 2019. ISSN 1367-4803. doi:10.1093/bioinformatics/bty1006. URL 10.1093/bioinformatics/bty1006.

[18] Jaspreet Singh, Thomas Litfin, Kuldip Paliwal, Jaswinder Singh, Anil Kumar Hanumanthappa, and Yaoqi Zhou. SPOT-1D-Single: improving the single-sequence-based prediction of protein secondary structure, backbone angles, solvent accessibility and half-sphere exposures using a large training set and ensembled deep learning. Bioinformatics, 37(20):3464–3472, October 2021. ISSN 1367-4803. doi:10.1093/bioinformatics/btab316. URL 10.1093/bioinformatics/btab316.

[19] Jaspreet Singh, Kuldip Paliwal, Thomas Litfin, Jaswinder Singh, and Yaoqi Zhou. Reaching alignment-profile-based accuracy in predicting protein secondary and tertiary structural properties without alignment. Scientific Reports, 12(1):7607, May 2022. ISSN 2045-2322. doi:10.1038/s41598-022-11684-w. URL https://www.nature.com/articles/s41598-022-11684-w.

[20] Burkhard Rost, Chris Sander, and Reinhard Schneider. PHD-an automatic mail server for protein secondary structure prediction. Bioinformatics, 10(1):53–60, February 1994. ISSN 1367-4803. doi:10.1093/bioinformatics/10.1.53. URL 10.1093/bioinformatics/10.1.53.

[21] C. Geourjon and G. Deléage. SOPMA: significant improvements in protein secondary structure prediction by consensus prediction from multiple alignments. Bioinformatics, 11(6):681–684, December 1995. ISSN 1367-4803. doi:10.1093/bioinformatics/11.6.681. URL 10.1093/bioinformatics/11.6.681.

[22] Eshel Faraggi, Tuo Zhang, Yuedong Yang, Lukasz Kurgan, and Yaoqi Zhou. SPINE X: Improving protein secondary structure prediction by multistep learning coupled with prediction of solvent accessible surface area and backbone torsion angles. Journal of Computational Chemistry, 33(3):259–267, 2012. ISSN 1096-987X. doi:10.1002/jcc.21968. URL https://onlinelibrary.wiley.com/doi/abs/10.1002/jcc.21968. _eprint: https://onlinelibrary.wiley.com/doi/pdf/10.1002/jcc.21968.

[23] Francesco Bettella, Dawid Rasinski, and Ernst Walter Knapp. Protein Secondary Structure Prediction with SPARROW. Journal of Chemical Information and Modeling, 52(2):545–556, February 2012. ISSN 1549-9596. doi:10.1021/ci200321u. URL 10.1021/ci200321u. Publisher: American Chemical Society.

[24] Alexey Drozdetskiy, Christian Cole, James Procter, and Geoffrey J. Barton. JPred4: a protein secondary structure prediction server. Nucleic Acids Research, 43(W1):W389–W394, July 2015. ISSN 0305-1048. doi:10.1093/nar/gkv332. URL 10.1093/nar/gkv332.

[25] Rhys Heffernan, Yuedong Yang, Kuldip Paliwal, and Yaoqi Zhou. Capturing non-local interactions by long short-term memory bidirectional recurrent neural networks for improving prediction of protein secondary structure, backbone angles, contact numbers and solvent accessibility. Bioinformatics, 33(18):2842–2849, September 2017. ISSN 1367-4803. doi:10.1093/bioinformatics/btx218. URL 10.1093/bioinformatics/btx218.

[26] Rhys Heffernan, Kuldip Paliwal, James Lyons, Jaswinder Singh, Yuedong Yang, and Yaoqi Zhou. Single-sequence-based prediction of protein secondary structures and solvent accessibility by deep whole-sequence learning. Journal of Computational Chemistry, 39(26):2210–2216, 2018. ISSN 1096-987X. doi:10.1002/jcc.25534. URL https://onlinelibrary.wiley.com/doi/abs/10.1002/jcc.25534. _eprint: https://onlinelibrary.wiley.com/doi/pdf/10.1002/jcc.25534.

[27] Minkyung Baek, Frank DiMaio, Ivan Anishchenko, Justas Dauparas, Sergey Ovchinnikov, Gyu Rie Lee, Jue Wang, Qian Cong, Lisa N. Kinch, R. Dustin Schaeffer, Claudia Millán, Hahnbeom Park, Carson Adams, Caleb R. Glassman, Andy DeGiovanni, Jose H. Pereira, Andria V. Rodrigues, Alberdina A. van Dijk, Ana C. Ebrecht, Diederik J. Opperman, Theo Sagmeister, Christoph Buhlheller, Tea Pavkov-Keller, Manoj K. Rathinaswamy, Udit Dalwadi, Calvin K. Yip, John E. Burke, K. Christopher Garcia, Nick V. Grishin, Paul D. Adams, Randy J. Read, and David Baker. Accurate prediction of protein structures and interactions using a three-track neural network. Science, 373(6557):871–876, August 2021. doi:10.1126/science.abj8754. URL https://www.science.org/doi/full/10.1126/science.abj8754. Publisher: American Association for the Advancement of Science.

[28] Milot Mirdita, Konstantin Schütze, Yoshitaka Moriwaki, Lim Heo, Sergey Ovchinnikov, and Martin Steinegger. ColabFold: making protein folding accessible to all. Nature Methods, 19(6):679–682, June 2022. ISSN 1548-7105. doi:10.1038/s41592-022-01488-1. URL https://www.nature.com/articles/s41592-022-01488-1. Number: 6 Publisher: Nature Publishing Group.

[29] Zeming Lin, Halil Akin, Roshan Rao, Brian Hie, Zhongkai Zhu, Wenting Lu, Nikita Smetanin, Robert Verkuil, Ori Kabeli, Yaniv Shmueli, Allan dos Santos Costa, Maryam Fazel-Zarandi, Tom Sercu, Salvatore Candido, and Alexander Rives. Evolutionary-scale prediction of atomic-level protein structure with a language model. Science, 379(6637):1123–1130, March 2023. doi:10.1126/science.ade2574. URL https://www.science.org/doi/10.1126/science.ade2574. Publisher: American Association for the Advancement of Science.

[30] Ruidong Wu, Fan Ding, Rui Wang, Rui Shen, Xiwen Zhang, Shitong Luo, Chenpeng Su, Zuofan Wu, Qi Xie, Bonnie Berger, Jianzhu Ma, and Jian Peng. High-resolution de novo structure prediction from primary sequence, July 2022. URL https://www.biorxiv.org/content/10.1101/2022.07.21.500999v1. Pages: 2022.07.21.500999 Section: New Results.

[31] Jennifer M. Michaud, Ali Madani, and James S. Fraser. A language model beats alphafold2 on orphans. Nature Biotechnology, 40(11):1576–1577, November 2022. ISSN 1546-1696. doi:10.1038/s41587-022-01466-0. URL https://www.nature.com/articles/s41587-022-01466-0. Number: 11 Publisher: Nature Publishing Group.

[32] Tomasz Smolarczyk, Irena Roterman-Konieczna, and Katarzyna Stapor. Protein Secondary Structure Prediction: A Review of Progress and Directions. Current Bioinformatics, 15(2):90–107, February 2020. doi:10.2174/1574893614666191017104639.

[33] Qian Jiang, Xin Jin, Shin-Jye Lee, and Shaowen Yao. Protein secondary structure prediction: A survey of the state of the art. Journal of Molecular Graphics and Modelling, 76:379–402, September 2017. ISSN 1093-3263. doi:10.1016/j.jmgm.2017.07.015. URL https://www.sciencedirect.com/science/article/pii/S1093326317304217.

[34] Supriyo Chakraborty, Richard Tomsett, Ramya Raghavendra, Daniel Harborne, Moustafa Alzantot, Federico Cerutti, Mani Srivastava, Alun Preece, Simon Julier, Raghuveer M. Rao, Troy D. Kelley, Dave Braines, Murat Sensoy, Christopher J. Willis, and Prudhvi Gurram. Interpretability of deep learning models: A survey of results. In 2017 IEEE SmartWorld, Ubiquitous Intelligence & Computing, Advanced & Trusted Computed, Scalable Computing & Communications, Cloud & Big Data Computing, Internet of People and Smart City Innovation (SmartWorld/SCALCOM/UIC/ATC/CBDCom/IOP/SCI), pages 1–6, August 2017. doi:10.1109/UIC-ATC.2017.8397411.

[35] Raul I. Perez Martell, Alison Ziesel, Hosna Jabbari, and Ulrike Stege. Supervised promoter recognition: a benchmark framework. BMC Bioinformatics, 23(1):118, April 2022. ISSN 1471-2105. doi:10.1186/s12859-022-04647-5. URL 10.1186/s12859-022-04647-5.

[36] Ričards Marcinkevičs and Julia E. Vogt. Interpretable and explainable machine learning: A methods-centric overview with concrete examples. WIREs Data Mining and Knowledge Discovery, 13(3):e1493, 2023. ISSN 1942-4795. doi:10.1002/widm.1493. URL https://onlinelibrary.wiley.com/doi/abs/10.1002/widm.1493. _eprint: https://onlinelibrary.wiley.com/doi/pdf/10.1002/widm.1493.

[37] Gwen R. Buel and Kylie J. Walters. Can AlphaFold2 predict the impact of missense mutations on structure? Nature Structural & Molecular Biology, 29(1):1–2, January 2022. ISSN 1545-9985. doi:10.1038/s41594-021-00714-2. URL https://www.nature.com/articles/s41594-021-00714-2. Number: 1 Publisher: Nature Publishing Group.

[38] Iqbal H. Sarker. Machine Learning: Algorithms, Real-World Applications and Research Directions. SN Computer Science, 2(3):160, March 2021. ISSN 2661-8907. doi:10.1007/s42979-021-00592-x. URL 10.1007/s42979-021-00592-x.

[39] J. R. Quinlan. Induction of decision trees. Machine Learning, 1(1):81–106, March 1986. ISSN 1573-0565. doi:10.1007/BF00116251. URL 10.1007/BF00116251.

[40] Leo Breiman. Random Forests. Machine Learning, 45(1):5–32, October 2001. ISSN 1573-0565. doi:10.1023/A:1010933404324. URL 10.1023/A:1010933404324.

[41] Pierre Geurts, Damien Ernst, and Louis Wehenkel. Extremely randomized trees. Machine Learning, 63(1): 3–42, April 2006. ISSN 1573-0565. doi:10.1007/s10994-006-6226-1. URL 10.1007/s10994-006-6226-1.

[42] Cullen Schaffer. When does overfitting decrease prediction accuracy in induced decision trees and rule sets? In Yves Kodratoff, editor, Machine Learning — EWSL-91, pages 192–205, Berlin, Heidelberg, 1991. Springer. ISBN 978-3-540-46308-5. doi:10.1007/BFb0017014.

[43] Krishna Teja Chitty-Venkata, Murali Emani, Venkatram Vishwanath, and Arun K. Somani. Neural Architecture Search for Transformers: A Survey. IEEE Access, 10:108374–108412, 2022. ISSN 2169-3536. doi:10.1109/ACCESS.2022.3212767. URL https://ieeexplore.ieee.org/abstract/document/9913476. Conference Name: IEEE Access.

[44] Frank Rosenblatt. The perceptron: a probabilistic model for information storage and organization in the brain. Psychological review, 65(6):386, 1958.

[45] Yann LeCun, Bernhard Boser, John Denker, Donnie Henderson, R. Howard, Wayne Hubbard, and Lawrence Jackel. Handwritten Digit Recognition with a Back-Propagation Network. In Advances in Neural Information Processing Systems, volume 2. Morgan-Kaufmann, 1989. URL https://proceedings.neurips.cc/paper/1989/hash/53c3bce66e43be4f209556518c2fcb54-Abstract.html.

[46] Alex Graves. Long Short-Term Memory. In Alex Graves, editor, Supervised Sequence Labelling with Recurrent Neural Networks, pages 37–45. Springer, Berlin, Heidelberg, 2012. ISBN 978-3-642-24797-2. doi:10.1007/978-3-642-24797-2_4. URL 10.1007/978-3-642-24797-2_4.

[47] Ashish Vaswani, Noam Shazeer, Niki Parmar, Jakob Uszkoreit, Llion Jones, Aidan N Gomez, Łukasz Kaiser, and Illia Polosukhin. Attention is All you Need. In Advances in Neural Information Processing Systems, volume 30. Curran Associates, Inc., 2017. URL https://proceedings.neurips.cc/paper_files/paper/2017/hash/3f5ee243547dee91fbd053c1c4a845aa-Abstract.html.

[48] J. Richard Landis and Gary G. Koch. A One-Way Components of Variance Model for Categorical Data. Biometrics, 33(4):671–679, 1977. ISSN 0006-341X. doi:10.2307/2529465. URL https://www.jstor.org/stable/2529465. Publisher: International Biometric Society.

[49] C. E. Shannon. A mathematical theory of communication. The Bell System Technical Journal, 27(3):379–423, July 1948. ISSN 0005-8580. doi:10.1002/j.1538-7305.1948.tb01338.x. URL https://ieeexplore.ieee.org/abstract/document/6773024. Conference Name: The Bell System Technical Journal.

[50] Paul D. Adams, Pavel V. Afonine, Kumaran Baskaran, Helen M. Berman, John Berrisford, Gerard Bricogne, David G. Brown, Stephen K. Burley, Minyu Chen, Zukang Feng, Claus Flensburg, Aleksandras Gutmanas, Jeffrey C. Hoch, Yasuyo Ikegawa, Yumiko Kengaku, Eugene Krissinel, Genji Kurisu, Yuhe Liang, Dorothee Liebschner, Lora Mak, John L. Markley, Nigel W. Moriarty, Garib N. Murshudov, Martin Noble, Ezra Peisach, Irina Persikova, Billy K. Poon, Oleg V. Sobolev, Eldon L. Ulrich, Sameer Velankar, Clemens Vonrhein, John Westbrook, Marcin Wojdyr, Masashi Yokochi, and Jasmine Y. Young. Announcing mandatory submission of PDBx/mmCIF format files for crystallographic depositions to the Protein Data Bank (PDB). Acta Crystallographica. Section D, Structural Biology, 75(Pt 4):451–454, April 2019. ISSN 2059-7983. doi:10.1107/S2059798319004522. URL https://www.ncbi.nlm.nih.gov/pmc/articles/PMC6465986/.

[51] Limin Fu, Beifang Niu, Zhengwei Zhu, Sitao Wu, and Weizhong Li. CD-HIT: accelerated for clustering the next-generation sequencing data. Bioinformatics, 28(23):3150–3152, December 2012. ISSN 1367-4803. doi:10.1093/bioinformatics/bts565. URL 10.1093/bioinformatics/bts565.

[52] Fabian Sievers and Desmond G. Higgins. Clustal Omega. Current Protocols in Bioinformatics, 48(1):3.13.1–3.13.16, 2014. ISSN 1934-340X. doi:10.1002/0471250953.bi0313s48. URL https://onlinelibrary.wiley.com/doi/abs/10.1002/0471250953.bi0313s48. _eprint: https://onlinelibrary.wiley.com/doi/pdf/10.1002/0471250953.bi0313s48.

[53] James A. Cuff and Geoffrey J. Barton. Evaluation and improvement of multiple sequence methods for protein secondary structure prediction. Proteins: Structure, Function, and Bioinformatics, 34(4):508–519, 1999. ISSN 1097-0134. doi:10.1002/(SICI)1097-0134(19990301)34:4<508::AID-PROT10>3.0.CO;2-4. URL https://onlinelibrary.wiley.com/doi/abs/10.1002/%28SICI%291097-0134%2819990301%2934%3A4%3C508%3A%3AAID-PROT10%3E3.0.CO%3B2-4. _eprint: https://onlinelibrary.wiley.com/doi/pdf/10.1002/%28SICI%291097-0134%2819990301%2934%3A4%3C508%3A%3AAID-PROT10%3E3.0.CO%3B2-4.

[54] Andriy Kryshtafovych, Torsten Schwede, Maya Topf, Krzysztof Fidelis, and John Moult. Critical assessment of methods of protein structure prediction (CASP)—Round XV. Proteins: Structure, Function, and Bioinformatics, 91(12):1539–1549, 2023. ISSN 1097-0134. doi:10.1002/prot.26617. URL https://onlinelibrary.wiley.com/doi/abs/10.1002/prot.26617. _eprint: https://onlinelibrary.wiley.com/doi/pdf/10.1002/prot.26617.

[55] Fabian Pedregosa, Gaël Varoquaux, Alexandre Gramfort, Vincent Michel, Bertrand Thirion, Olivier Grisel, Mathieu Blondel, Peter Prettenhofer, Ron Weiss, Vincent Dubourg, Jake Vanderplas, Alexandre Passos, David Cournapeau, Matthieu Brucher, Matthieu Perrot, and Édouard Duchesnay. Scikit-learn: Machine Learning in Python. J. Mach. Learn. Res., 12(null):2825–2830, November 2011. ISSN 1532-4435.

[56] D. J. Newman. Simple Analytic Proof of the Prime Number Theorem. The American Mathematical Monthly, 87(9):693–696, November 1980. ISSN 0002-9890. doi:10.1080/00029890.1980.11995126. URL 10.1080/00029890.1980.11995126. Publisher: Taylor & Francis _eprint: 10.1080/00029890.1980.11995126.

[57] Gürol Canbek, Seref Sagiroglu, Tugba Taskaya Temizel, and Nazife Baykal. Binary classification performance measures/metrics: A comprehensive visualized roadmap to gain new insights. In 2017 International Conference on Computer Science and Engineering (UBMK), pages 821–826, October 2017. doi:10.1109/UBMK.2017.8093539. URL https://ieeexplore.ieee.org/document/8093539/figures#figures.

[58] Fu V Song, Jiaqi Su, Sixing Huang, Neng Zhang, Kaiyue Li, Ming Ni, and Maofu Liao. DeepSS2GO: protein function prediction from secondary structure. Briefings in Bioinformatics, 25(3):bbae196, May 2024. ISSN 1477-4054. doi:10.1093/bib/bbae196. URL 10.1093/bib/bbae196.

