## Supplementary materials for "Missense mutation knowledge can decrease prediction inaccuracies on protein secondary structure"

#### 1. MUTATION EXTRACTION

Finding mutations from an a series of aligned sequences and assigning a consensus or mutation value to a sequence is non-trivial. To find mutations in our preprocessed data, we created a similar method from Weblogo [1] which use information theory to provide the significance of each mutation. The height of each mutation in the logo is characterized by the frequency of the amino acid at a specific position ($p_i$) and subsequently through Shannon entropy,

$$S = -\sum_i p_i \log_2 p_i \tag{S1}$$

Equation S1 ranges from $[0, 1]$ with a domain of frequencies of $[0, 1]$ and has a characteristic bell shaped curve with a maximum on 0.5. Therefore, values for frequencies that are equally spaced apart from the maximum value give the same result. Unfortunately, we require a method that can distinguish between such values since the consensus frequency might be equally spaced apart from the mutation frequency (e.g. 0.25 and 0.75), but clearly one is more frequent than the other. Therefore we decided to modify the equation as follows,

$$S = -\sum_i p_i \log_2 (1 - p_i) \tag{S2}$$

This was done to obtain mutation positions easily since positions without mutations will return 0 or Infinity. Infinity values are subsequently turned to 0. Any non-zero value will be valid for a mutation or consensus amino acid in a position where a mutation has occurred. By using sparse matrices, it is then simple to systematically find these non-zero values and assign its sequence either a consensus (maximum value) or mutation (others) category for a location in the sequence.

#### 2. DSSP DISCREPANCIES

We used DSSP to assign secondary structures to the protein MMCIF' structure data. The macromolecular crystallographic information file (mmCIF) format was selected because the PDB has required it for all new submissions since 2019 [2]. As a result, structures submitted to the PDB after 2019 are no longer available in the older PDB' format, rendering older software incompatible with recent data. DSSP has been updated to process the mmCIF format and has been extensively tested by the community.

DSSP's algorithm for assigning secondary structures to mmCIF-formatted input has evolved from its original version, which processed 'PDB' files. The main differences include both the file formatting and the secondary structure classification, which varies from the traditional 8 classes, as shown in Table S1.

When producing mmCIF output, DSSP expands the input file by rewriting it and appending the secondary structure information. However, this rewriting process can introduce formatting errors, particularly when the input file contains quotations with the character **'**. To address these issues, we provide a script that properly re-formats DSSP's mmCIF output.

As noted earlier and shown in Table S1, the mmCIF format classifies both the $\beta$-bridge and Strand under the same STRN class, making it difficult to distinguish between them. Kabash and Sander, in their original DSSP publication, refer to $\beta$-bridge as an "isolated bridge", which is formed by a single hydrogen bond similar to those found in a $\beta$-sheet (DSSP's Strand class).

According to Kabash and Sander's description of secondary structures, an isolated bridge is part of the repeating hydrogen-bonding patterns "turn" and "bridge." Repeating turns form "helices," repeating bridges form "ladders," and connected ladders form "sheets." Based on this understanding, we convert the mmCIF output back to the original classes used in the 'PDB' format, which are recognized by all secondary structure prediction tools. Specifically, when we encounter an isolated STRN, we convert it to its corresponding $\beta$-bridge (**B**) class.

It's also important to note that the PolyProline (PPII) class is a newer addition in the mmCIF format. However, since prediction methods rely on the Q8 assignment, the PPII class is not used in our analysis. Additionally, the OTHER class does not appear in the output, as DSSP only

| DSSP Class (Q8) | PDB Class | mmCIF Class | Description |
|---|---|---|---|
| B | B | STRN | $\beta$-bridge |
| C | ' '(space) | OTHER | Loop or Coil |
| E | E | STRN | Strand |
| G | G | HELX_RH_3T_P | 3-10 helix |
| H | H | HELX_RH_AL_P | $\alpha$-helix |
| I | I | HELX_RH_PI_P | $\pi$-helix |
| S | S | BEND | Bend |
| T | T | TURN_TY1_P | Turn |
| | P | HELX_RH_PP_P | PPII-helix |

**Table S1.** DSSP class conversion by input format. Note that classes $\beta$-bridge and Strand are indistinguishable by the mmCIF DSSP algorithm.

generates proper secondary structural classes. The OTHER class can be inferred when a residue lacks a structural classification.

By applying these conversions, we can accurately assess mmCIF-formatted structure files in terms of the original Q8 classification scheme.

#### 3. MUT2DENS MODEL DETAILS

The model Mut2dens consists of extremely randomized trees trained using scikit-learn[1] with default hyperparameter values. We utilized the default hyper-parameters for training the model. The input consists of nominal windowed data concatenated with a window length of 7, or window side length of 3. We utilized this window length as it was the best window length for top predictors. Therefore, the input data can be viewed as a vector of length $m = 7 \cdot p$, where $p$ is the amount of predictors utilized. The best model during testing utilized an input combination of SSPro8, ColabFold, and ESMFold. Thus, the final Mut2dens model makes use of three predictors ($p = 3$) and a window size of 21 ($m = 21$) for Mut2dens final model. This model slightly sacrified mean performance to increase the performance on inaccurate predictions.

Training the model takes less than 30 seconds using 8 cores from an Intel Xeon 2600 family CPU, while inference takes less than 3 seconds. Therefore, the majority of the time taken when utilizing this refinement procedure is spent obtaining predictions from the desired predictors. The time efficiency of the predictors is given below.

#### 4. PREDICTORS COMPUTATIONAL PERFORMANCE

To run the predictors, we utilize cloud resources which contain common server infrastructure CPU resources, such as the Intel Xeon 2600 family of processors. We utilized 16-core CPUs as a greater number of cores did not seem to improve performance significantly. When utilizing GPU resources, we utilized a single Nvidia V100. Per protein, with an average length of 300 amino acids, the average total time taken for each predictor is given in Table S2

It is important to note that most of the time taken to predict proteins from these predictors come from their use of multiple sequence alignment or physics-based atomic relaxation techniques. These procedures can average around 60% to 80% of the predictor's time taken during prediction which we include to account for all processing done after a protein sequence is inputted into the predictor.

[1] https://scikit-learn.org/stable/modules/generated/sklearn.ensemble.ExtraTreesClassifier.html

| Predictor | Average time taken |
|---|---|
| AlphaFold2 | $\sim$ 2 hours$^+$ |
| ColabFold | $\sim$ 30 minutes$^+$ |
| SSPro8 | $\sim$ 3 hours$^+$ |
| ESMFold | $\sim$ 5 minutes |
| SPOT1D-LM | $\sim$ 5 minutes |
| SPOT1D | $\sim$ 4 hours$^+$ |
| SPOT1D-Single | $\sim$ 5 minutes |
| RGN2 | $\sim$ 5 minutes* |
| Raptor-X Property | $\sim$ 1 minute |

**Table S2.** * Time taken using a truncated amount of atomic relaxation. Without truncation, time could exceed 2 hours. $^+$ utilizes MSA procedure.

#### 5. MACHINE LEARNING MODELS

In this section we describe the architectural and hyperparameter details of the multiple machine learning models we investigated. The models investigated include tree-type and neural-type models.

##### A. Tree-type model details

All tree-type models were created as classifiers from sci-kit learn. Their hyperparameters follow their default values, which are shown in Table S3.

| Property | Decision Tree | Random Forest | Extra Trees |
|---|---|---|---|
| Split criterion | Gini Impurity | Gini Impurity | Gini Impurity |
| Trees | 1 | 100 | 100 |
| Minimum split samples | 2 | 2 | 2 |
| Minimum leaf samples | 1 | 1 | 1 |
| Features considered | All ($M$) | $\sqrt{M}$ | $\sqrt{M}$ |
| Bootstrap samples | No | No | No |
| Pruning | No | No | No |

**Table S3.** All tree-type models were created as similar as possible. For the decision tree, all features had to be considered as only a single tree is created. Bootstrapping and Pruning were not utilized to avoid removing any possible context from all predictors. Overfitting was only an issue after window lengths increased. If greater window lengths are required, utilizing these techniques could potentially alleviated overfitting.

##### B. Neural-type model details

All neural-type architectures were developed using PyTorch. The diagrams and details of the different architectures are given below.

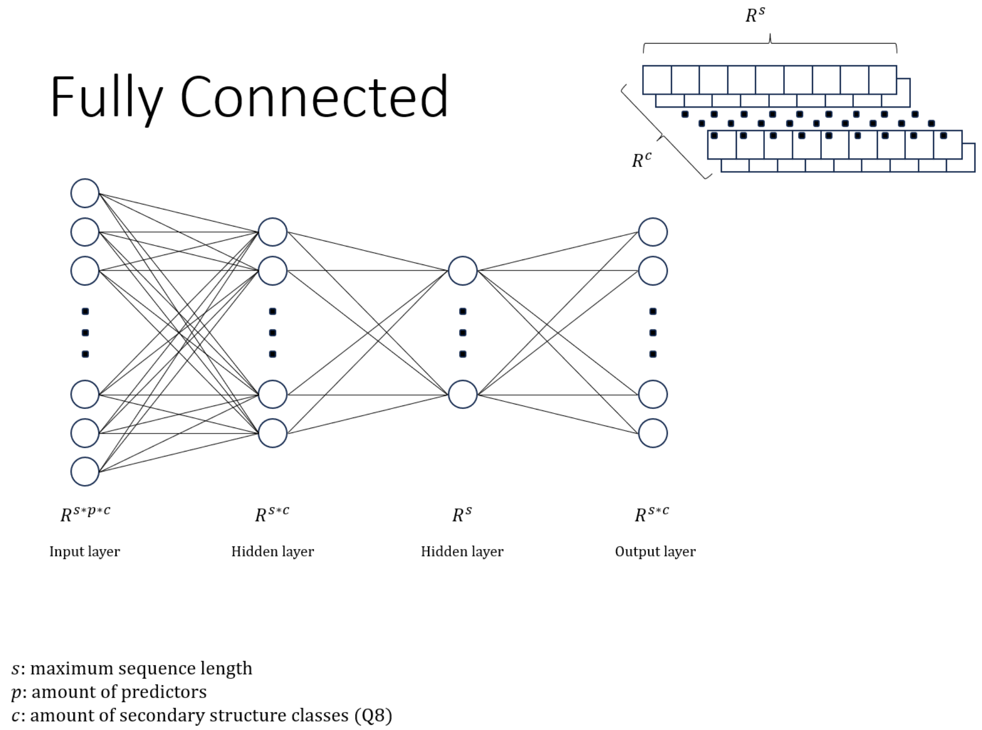


**Fig. S1. Fully-connected architecture.** Each layer of the network consists of a linear layer that connects all neurons to the next layer followed by batch normalization, a rectified linear unit (ReLU) activation function, and a 20% neuron dropout probability.

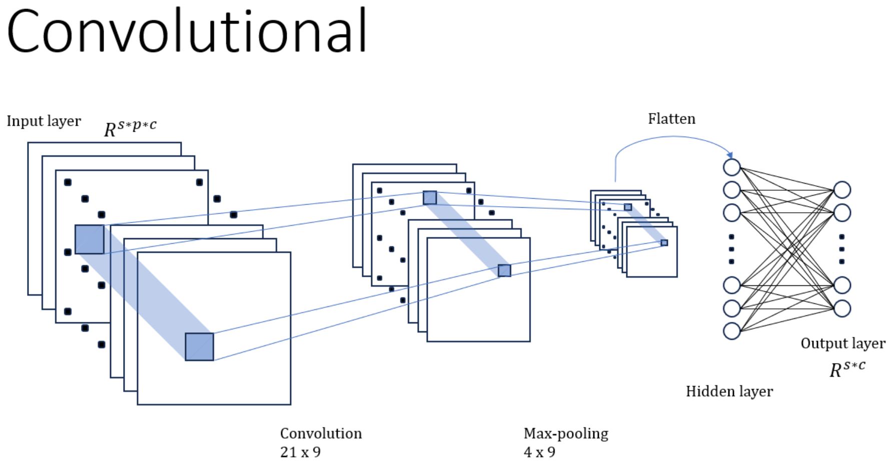


**Fig. S2. Convolutional architecture.** Each layer of the network consists of a convolutional layer with a halving amount of channels starting from 64. Each layer also contains batch normalization, a ReLU activation function, and a 20% neuron dropout probability. The convolutions are then flattened into a vector and passed through a final linear layer as in the fully-connected architecture.

### Recurrent

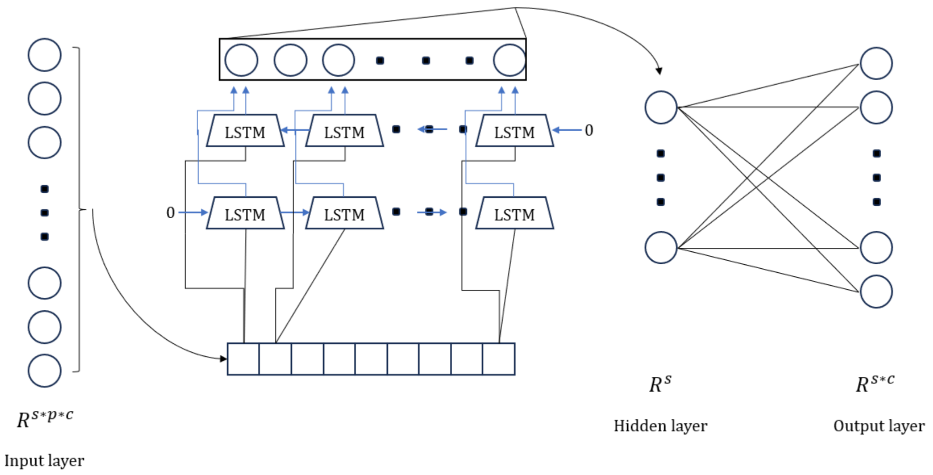


**Fig. S3. Recurrent architecture.** It consists of two long short-term memory (LSTM) layers that process the input in opposite directions, also known as a bidirectional LSTM. One processes the input from start to end, while the other from end to start. The LSTM layers also contain a 10% probability of neuron dropout. The LSTM layers' output is combined and passed through a final linear layer as in the previous architectures.

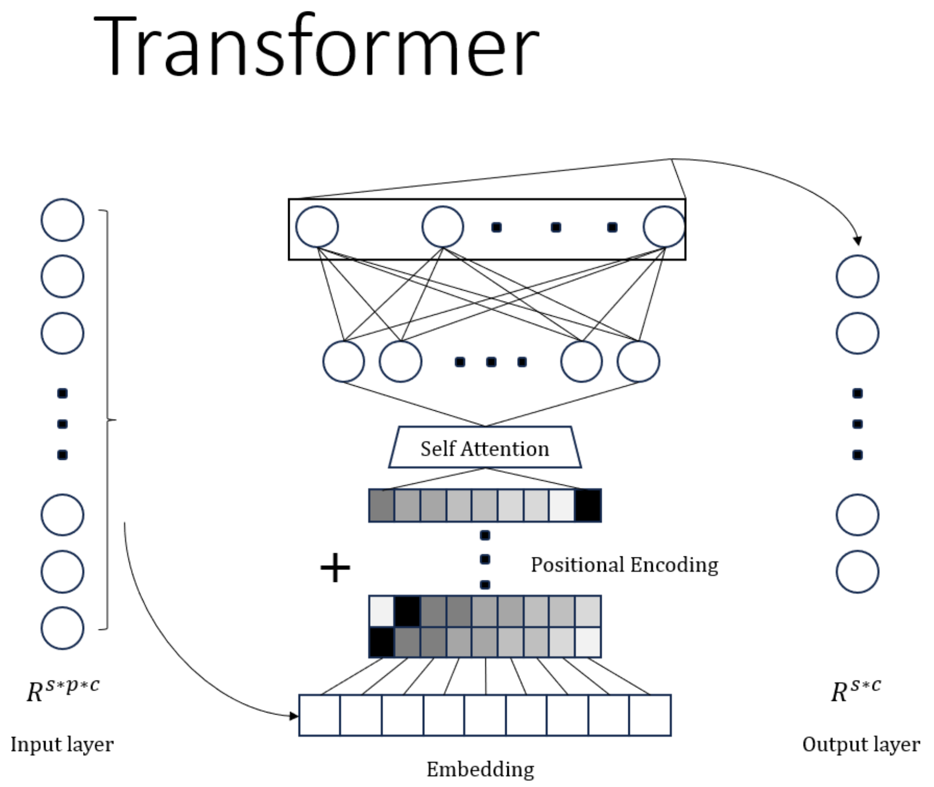


**Fig. S4. Transformer architecture.** It first embeds the input into vectors to be processed by positional encoding. This is then passed to two self-attention layers with 50% neuron dropout probability. The attention layer outputs are passed to intermediary fully-connected linear layers and then their outputs are combined and passed to a final linear layer as in the previous architectures.

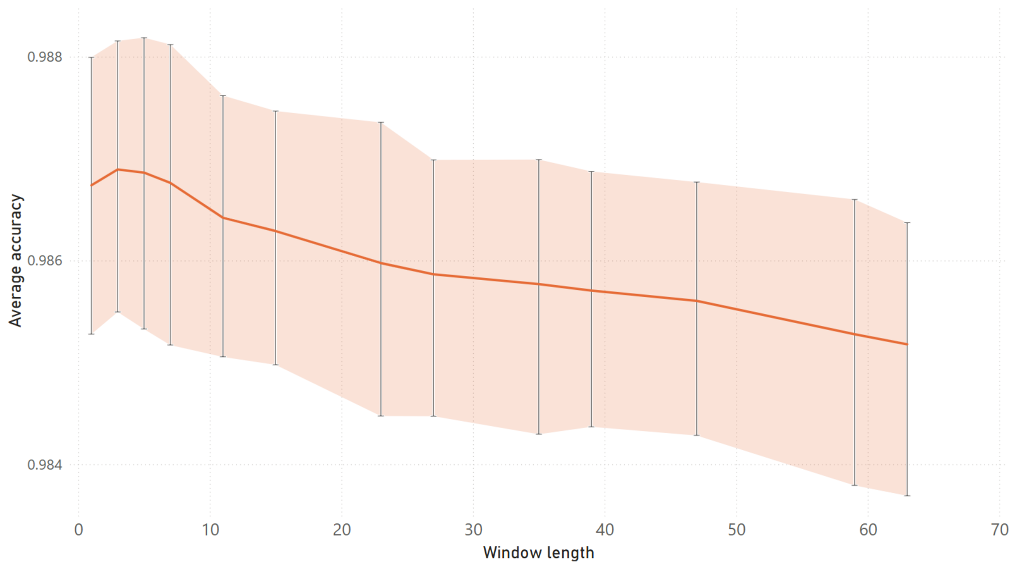


**Fig. S5. Training accuracy of top-performing predictors for different window lengths.** Diagram showing performance details for top-performing methods where an ExtraTree model has lower accuracy as the window length gets longer. This decrease might seem minuscule but it transfers remarkably well to test datasets, like CASP15. Within low window lengths, the performance of models increase until about a window length of 7. Afterwards, performance deteriorates quickly for unseen data. The reason for this is likely to be from overfitting the data with longer window lengths as the limited dataset provide a decreasing number of data points as the window size increases.

#### REFERENCES

1. Crooks GE, Hon G, Chandonia JM, Brenner SE. WebLogo: A Sequence Logo Generator. Genome Research. 2004;14(6):1188–1190. doi:10.1101/gr.849004.
2. Adams PD, Afonine PV, Baskaran K, Berman HM, Berrisford J, Bricogne G, et al. Announcing mandatory submission of PDBx/mmCIF format files for crystallographic depositions to the Protein Data Bank (PDB). Acta Crystallographica Section D, Structural Biology. 2019;75(Pt 4):451–454. doi:10.1107/S2059798319004522.
